# DNA Sequence-Programmed Protein Coronas Determine Intracellular Fate and Proteostatic Stress of Carbon Nanotubes

**DOI:** 10.64898/2026.08.31.748348

**Authors:** Aceer Nadeem, Tianyu Xu, Jacob Miller, Delaney Kirk, Mijin Kim

**Affiliations:** School of Chemistry and Biochemistry, Georgia Institute of Technology, Atlanta, GA 30332, United States; School of Chemical and Biomolecular Engineering, Georgia Institute of Technology, Atlanta, GA 30332, United States; Parker H. Petit Institute for Bioengineering and Biosciences, Georgia Institute of Technology, Atlanta, GA 30332, USA

**Keywords:** single-walled carbon nanotubes, DNA surface functionalization, protein corona, intracellular trafficking, macrophage, near-infrared fluorescence, proteomics, precision nanotoxicology

## Abstract

Single-walled carbon nanotubes (SWCNTs) show promise for optical biosensing, imaging, and drug delivery, but turning them into safe, precision nanomedicine tools requires understanding how nanotube surface chemistry dictates recognition and processing by cells. Like other nanomaterials, carbon nanotubes acquire a biomolecular corona on contact with biological fluids, and corona identity is increasingly recognized as central to sensor performance and drug delivery efficacy. However, whether corona identity also governs the intracellular fate of carbon nanotubes remains largely unknown. Here, we show that the single-stranded DNA wrapping of (6,5)-enriched single-walled carbon nanotubes reprograms their protein corona, intracellular trafficking, and macrophage response. By profiling (AT)_15_, (GT)_15_, and (CT)_15_ wrapped SWCNTs, we show that the wrapping sequence programs both the protein corona and the resulting proteostatic stress on macrophages. Photoluminescence imaging and confocal Raman measurements reported that (AT)_15_ is internalized the most yet leaves the proteome and nanotube structure largely undisturbed, whereas (CT)_15_, taken up the least, undergoes the most aggressive intracellular degradation and drives the highest oxidative and proteostatic stress. Corona proteomics indicated that all three tested nanotubes form coronas with distinct functional identities that are responsible for divergent intracellular routes. Time-resolved intracellular proteomics combined with functional assays resolved how the host cell reorganizes its biomolecular complexity over time, including oxidative outputs, aside from a sequence-independent core response involving particle engagement, phagosomal sorting, and lysosomal processing. These findings provide mechanistic insight into nanomaterial-cell interactions and the wrapping sequence as a tunable, nucleotide-level design handle for controlling the intracellular fate of carbon nanomaterials, with potential implications for safe and effective nanomedicine platforms.

---

Carbon nanotubes (CNTs) can interface with biological systems at the molecular scale and have been explored for drug delivery^1^, real-time imaging^2,3^, and biochemical sensing^4,5^, but translating these into safe, clinically viable tools has proven difficult^6–9^. Rising concerns and limited information on their toxicological and environmental risks have led the International Chemical Secretariat correspondences and the EU Observatory for Nanomaterials to call for restricting the use of carbon nanotubes in the European Union, and ongoing CNT-related regulations and research by governmental agencies^10,11^. While nanotube toxicity as a function of structural descriptors such as length, charge, and aggregation states has been extensively investigated,^12–15^ structurally similar CNTs give inconsistent and at times contradictory toxicity results across studies^16–18^, suggesting that other properties of the nanotube surface not captured by these descriptors drive different biological outcomes^19,20^.

Biomolecular corona is a layer of biomolecules that assembles on a nanomaterial upon contact with biological fluid^21^, and the most critical molecular feature that carries this surface identity into biological systems. Corona composition governs receptor engagement^22,23^, endocytic routing^24^, immune recognition^25,26^, and intracellular trafficking^27^. On carbon nanotubes, we are beginning to understand how corona composition varies with surface chemistry.^28–30^ Because corona identity is shaped by more than bulk descriptors, carbon nanotubes indistinguishable by length, aggregation state, diameter, etc. can present entirely different biological interfaces^31^ and nanotoxicity^32–35^. However, systematic links between corona composition, intracellular trafficking, and downstream cellular and immune responses of carbon nanotubes, under controlled surface chemistry, remain a critical knowledge gap.

DNA-wrapped single-walled carbon nanotubes (DNA-SWCNTs) are well suited to close this gap because the wrapping sequence can be varied while keeping the coarse structural parameters constant, isolating surface chemistry as the sole variable. Single-stranded DNA adopts sequence-dependent conformations on the nanotube surface^14,36,37^, and reshapes the local interface, including its hydration shell^38,39^, hydrogen-bonding network^15,40,41^, charge distribution^42^, and corona composition. We therefore hypothesize that sequence-specific ssDNA wrapping influences intracellular trafficking and the macrophage response via protein corona engineering.

To test this hypothesis, we investigated (6,5) chirality-enriched SWCNTs wrapped with three sequences, (AT)_15_, (GT)_15_, and (CT)_15_, that differ in base composition and surface conformation but share an identical nanotube core and comparable net charge and defect density. Using RAW 264.7 macrophages as the cellular model, we combined protein-corona proteomics, time-resolved NIR hyperspectral imaging, confocal Raman microscopy, cell-based functional assays, and intracellular proteomics to follow each construct from corona formation through uptake to intracellular fate and cellular response. We found that the wrapping sequence sets the intracellular fate of DNA-SWCNTs independently of how much material a macrophage takes up. (AT)_15_-SWCNTs are internalized the most yet retain an essentially intact nanotube sidewall and never mount a significant oxidative response, whereas (CT)_15_-SWCNTs, taken up the least, undergo the most aggressive structural degradation, the largest cumulative proteome disruption, and a 7-fold rise in reactive oxygen species. (GT)_15_-SWCNTs showed minimal oxidative burden paired with a 10-fold rise in intracellular ATP, marking a metabolic response to nanotube exposure. These divergent trajectories track the degree of lysosomal engagement and map onto compositionally distinct plasma protein coronas recruited by each sequence, decoupling nanotoxicity from the length-, charge-, and aggregation-based descriptors that dominate current risk-assessment frameworks. By resolving corona identity, intracellular trafficking, and proteostatic output for three otherwise physicochemically matched constructs, we establish nucleotide sequence as a programmable, single-residue-level design variable for controlling how cells recognize, process, and tolerate carbon nanomaterials, providing a mechanistic foundation for engineering safer nanomedicine platforms.

## RESULTS

### (6,5)-Enriched DNA-SWCNTs Form Physicochemically Equivalent Dispersions

To isolate the wrapping sequence as a single variable, we first prepared three DNA-SWCNT constructs matched in chirality, charge, and defect density. (6,5)-SWCNTs were purified with aqueous two-phase extraction,^43^ followed by polymer rewrapping^44^ (**Figure 1A**). UV-vis-NIR absorption spectra and excitation-emission fluorescence maps indicate an enriched (6,5) fraction as high as 75%, with trace amounts of (6,4)– and (8,4)-SWCNTs (**Figures 1B**, Figures S1-S2, Tables S1-S2). Successful surfactant-to-DNA exchange was confirmed by the characteristic emission redshift from 985 nm to above 990 nm upon DNA wrapping (**Figure 1B**, Table S3)^45,46^.

**Figure 1.**
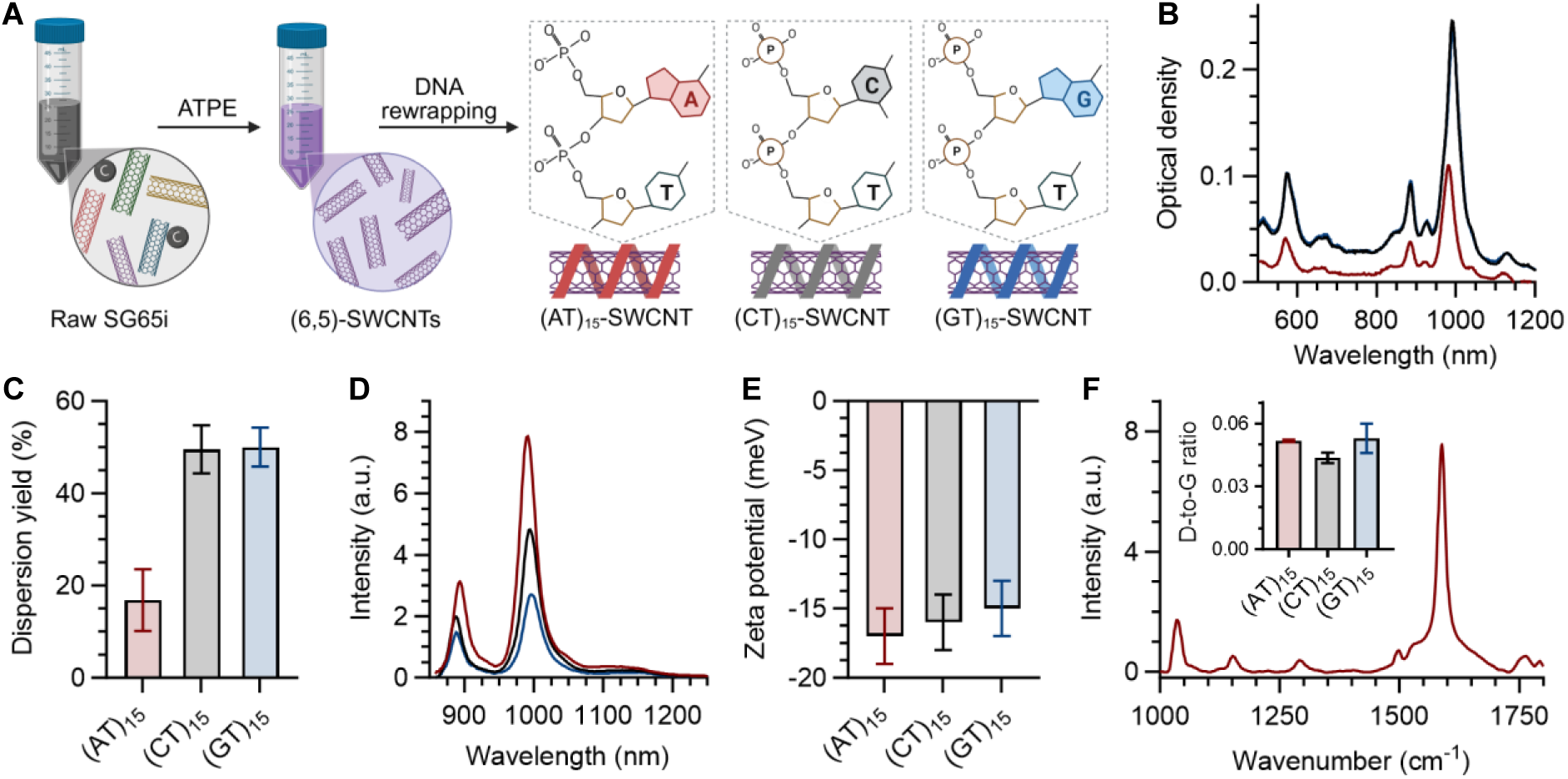
Three DNA-SWCNT constructs, physicochemically equivalent and differ only in wrapping sequence. (**A**) Preparation. Raw CoMoCAT SG65i SWCNTs were enriched for the (6,5) chirality by aqueous two-phase extraction (ATPE) and rewrapped with ssDNA. (**B**) UV-vis-NIR absorption spectra of the three DNA-SWCNT dispersions. Red: (AT)_15_-SWCNT. Black/gray: (CT)_15_-SWCNT. Blue: (GT)_15_-SWCNT. (**C**) Dispersion yield, defined as the mass fraction of SWCNTs recovered in the DNA-SWCNT supernatant relative to the starting stock. (**D**) NIR emission spectra at a matched SWCNT concentration of 1 mg L^-1^ under 575 nm excitation. (**E**) Zeta potential of DNA-SWCNT (5 mg/L in water). (**F**) Representative Raman spectrum of a (AT)_15_-SWCNT under 785 nm excitation. Inset, the D-to-G intensity ratio for the three constructs. Bars are mean ± s.d. of n = 3.

The three wrapping sequences, (AT)_15_, (CT)_15_, and (GT)_15_, produced clear differences in dispersion quality and decoupled dispersion efficiency from optical brightness. Dispersion yield, defined as the mass fraction of nanotubes recovered in the DNA-SWCNT supernatant relative to the starting SWCNT stock, was substantially lower for (AT)_15_ than the other two sequences (**Figure 1C**), yet at matched SWCNT concentration (1 mg L^-1^), (AT)_15_-SWCNTs showed the brightest per-mass NIR emission (**Figure 1D**). Zeta potential, Raman D-to-G ratio, and chirality distribution were statistically indistinguishable across all three DNA-SWCNT constructs (**Figures 1E-F**, Table S2), indicating that the nanotube core, charge, and defect density are comparable.

### DNA Sequence-Dependent Intracellular Uptake and Retention Dynamics

We exposed RAW 264.7 macrophages to each DNA-SWCNT construct at a concentration of 1mg L^-1^ and tracked NIR optical behavior and Raman signal over time (**Figure 2A**). All three ssDNA-SWCNT constructs preserved macrophage viability over 24 hours (Figure S3). At 1 hour, the fluorescence intensity of (AT)_15_ and (GT)_15_-SWCNTs was comparable in cells, while (CT)_15_-SWCNTs already showed significantly lower fluorescence (**Figures 2B-C**, Figures S4-S5). Over 24 hours, fluorescence imaging showed that (AT)_15_-SWCNTs retain the large majority of their emission while (GT)_15_– and especially (CT)_15_-SWCNTs undergo a substantial decrease in fluorescence (∼75%).

**Figure 2.**
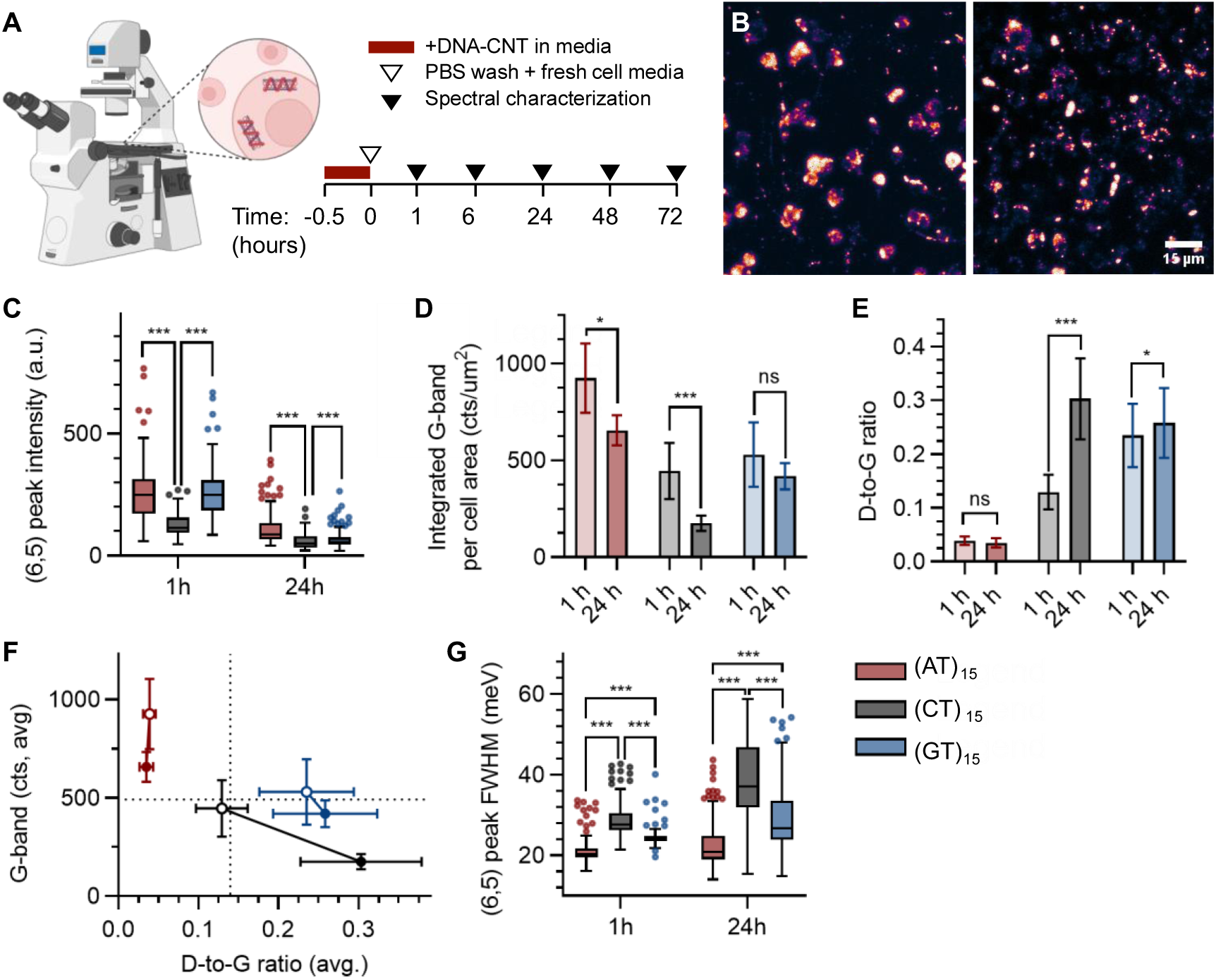
Sequence-dependent intracellular uptake, retention, structural fate, and NIR optical behavior of DNA-SWCNTs in macrophages. (**A**) Experimental workflow. RAW 264.7 macrophages were exposed to DNA-SWCNTs (1 mg L^-1^) in culture medium for 30 min, washed with PBS, and returned to fresh medium (t = 0). Cells were characterized by NIR hyperspectral fluorescence and confocal Raman spectroscopy at 1, 6, 24, 48, and 72 h. (**B**) Representative NIR fluorescence images of macrophages treated with (AT)_15_-SWCNTs at 1 h (left) and 24 h (right). Scale bar: 15 µm. (**C**) (6,5) emission peak intensity per cell at 1 and 24 h for each wrapping sequence. Boxes show the median and interquartile range (IQR), whiskers extend to 1.5x IQR (Tukey), and points beyond the whiskers are outliers. (**D**) Integrated Raman G-band intensity per cell area at 1 and 24 h. (**E**) Raman D-to-G band ratio at 1 and 24 h. (**F**) Joint trajectory of nanotube abundance and structural integrity, plotting average G-band intensity against average D-to-G ratio for each construct at 1 h (open) and 24 h (filled). Connecting arrows indicate temporal trajectory from 1h to 24h. Dotted reference lines are included as visual guides to aid comparison of relative changes in nanotube abundance and structural integrity across constructs. (**G**) Full width at half-maximum (FWHM) of the (6,5) emission peak at 1 and 24 h. Statistical significance: *p < 0.05, **p < 0.01, ***p < 0.001; ns, not significant. Red: (AT)_15_-SWCNT. Black/gray: (CT)_15_-SWCNT. Blue: (GT)_15_-SWCNT.

To track total nanotube quantity in cells, including both fluorescent and quenched species^47^, we probed the sp^2^-graphitic G-band (1,585 cm^-1^) of SWCNTs using confocal Raman spectroscopy as a structure-specific marker. The Raman G-band quantification of the fixed cells confirmed the uptake hierarchy at 1 hour of (AT)_15_ > (GT)_15_ > (CT)_15_ (**Figure 2D**). Over 24 hours, (AT)_15_– and (GT)_15_-SWCNTs retained 71% and 79% of their initial G-band signal, respectively, indicating that most of the internalized material remains within macrophages. In contrast, (CT)_15_-SWCNTs lost 40% of their initial G-band intensity due to rapid clearance or active exocytosis.

We next examined the Raman D-band-to-G-band ratio to assess oxidative damage on nanotube sidewalls (**Figure 2E**). The Raman D-band at 1,350 cm^-1^ arises from symmetry breaking of the sp^2^ graphitic lattice of carbon nanotubes, and thus the Raman D-to-G ratio is a relative measure of defect density^48–50^. The three constructs differed in both their starting defect ratios and trajectories through time. (AT)_15_-SWCNTs began with the lowest D-to-G ratio, comparable to the prior incubation (**Figure 1F**), and showed no statistically significant changes over 24 hours, with a slight decrease (–10.3%), indicating that the sidewall remained intact across intracellular residence. (CT)_15_-SWCNTs began at an intermediate value and rose sharply by 24 hours (+134.9%). (GT)_15_-SWCNTs showed the highest D-to-G ratio among the three constructs at 1 hour but increased only marginally through 24 hours (+9.8%). Defect accumulation in (GT)_15_-SWCNTs therefore occurred within the first hour rather than progressively through 24 hours over time, in contrast to what we observed with (CT)_15_-SWCNTs.

To jointly track nanotube abundance and structural integrity over time, G-band intensity is visualized against the D-to-G ratio for each construct (**Figure 2F**, Table S4). At 1 hour, (AT)_15_-SWCNTs sit in the high abundance, low defect quadrant, while (CT)_15_– and (GT)_15_-SWCNTs occupy relatively moderate and intermediate positions, respectively. Over 24 hours, (AT)_15_-SWCNTs undergo modest vertical displacement, declining in abundance while maintaining the defect density. (GT)_15_-SWCNT shows minimal displacement in either axis. (CT)_15_-SWCNTs simultaneously lose G-band intensity and accumulate structural defects.

The temporal evolution of fluorescence intensity and Raman signals indicates that the nanotube surface environment changes over the course of intracellular residence. Changes in the peak wavelength and full width at half-maximum (FWHM) of the (6,5) emission, both of which reflect the local dielectric^46,51^, aggregation states and population homogeneity^51^ support our observation (**Figure 2G**, Figure S5). (AT)_15_-SWCNTs display the narrowest FWHM at 1 h (21 meV) and maintain wavelength stability across the entire observation window, consistent with superior colloidal stability and residence in a uniform compartment with minimal perturbation of the nanotube surface. (CT)_15_-SWCNTs exhibited a large FWHM and broad distribution at 1 h, indicating substantial bundling from the earliest time point. FWHM distribution broadened further by 24 h due to dynamic remodeling of the interfacial environment on DNA-SWCNTs. The FWHM of (GT)_15_-SWCNTs began with an intermediate median (24 meV) and the narrowest distribution at 1 h, the FWHM redshifted progressively, and the distribution significantly broadened by 24 h, suggesting increasing heterogeneity and aggregation over time. These measurements show separable uptake magnitude and structural fate.

Extended spectral characterization over 72 h supports persistent trends observed at 24 h (Figures S6-S7). (AT)_15_-SWCNTs signal declined gradually while remaining spectrally stable, whereas (CT)_15_– and (GT)_15_-SWCNTs emission redshifted further and fell below the detection limit. Because the sequence-dependent behavior developed within the first 24 h and persisted thereafter, we analyzed the first 24 h, where the three constructs separate, for the remainder of the study.

### (CT)_15_ Wrapping Shapes the Most Compositionally Distinct Protein Corona

To resolve how the wrapping sequence shapes the biomolecular interface of the SWCNTs and how it can impact the intracellular fates, we characterized the protein corona recovered from DNA-SWCNTs by LC-MS/MS (**Figure 3A**). The log-transformed fold-change (log_2_FC) profiles (protein corona vs. full plasma) were compared across 241 retained plasma proteins (See Methods; Figure S8). Applying an enrichment cutoff of 1.5-fold change, 107 proteins were enriched in at least one corona, of which 45 were common to all three (**Figure 3B**). The three coronas recruited a near-identical number of enriched proteins, indicating that the wrapping sequence does not alter the overall breadth of protein adsorption. Pairwise correlation across the full corona dataset confirmed that the (CT)_15_ corona is the most compositionally distinct, and that the (AT)_15_ and (GT)_15_ coronas form the most similar pair (Figure S9).

**Figure 3.**
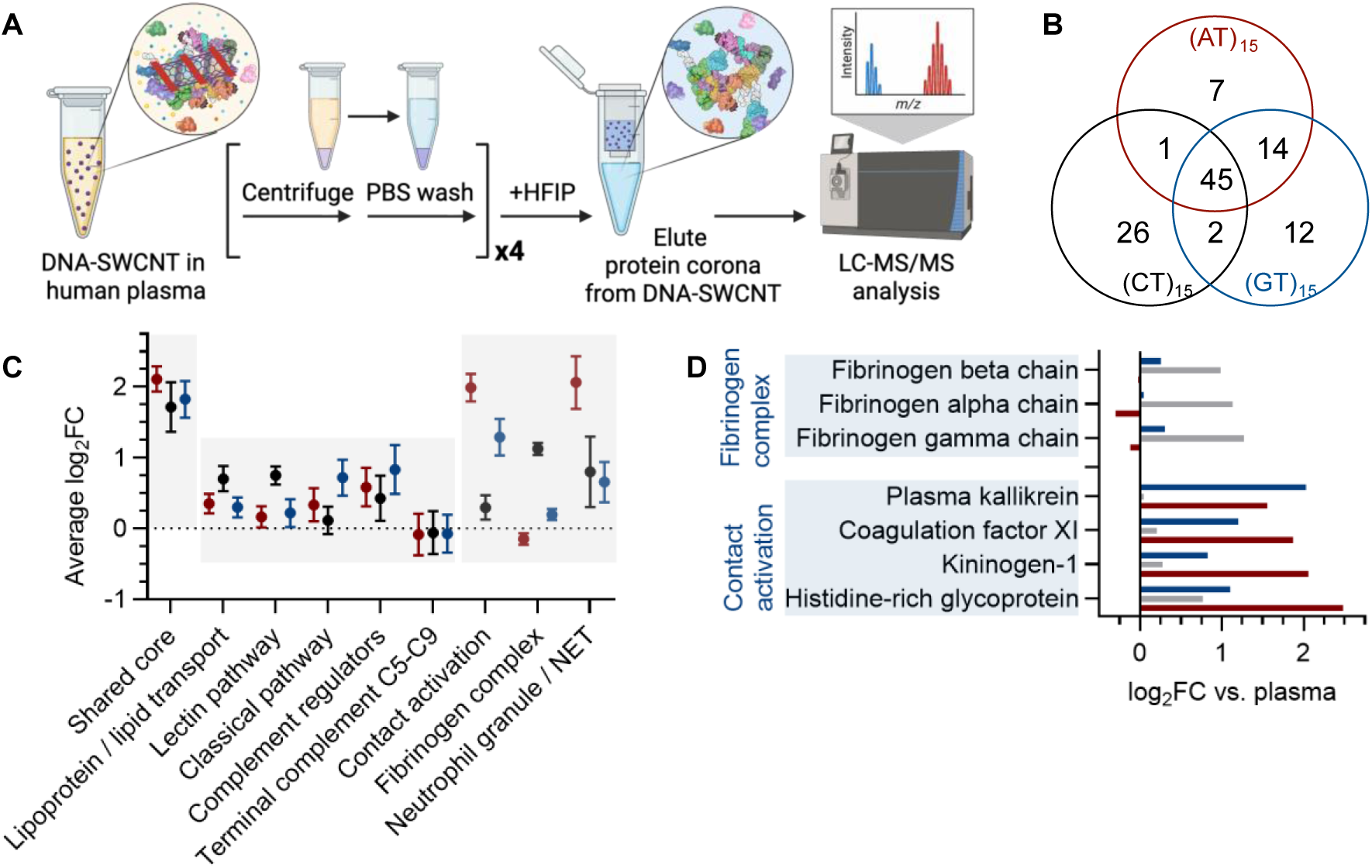
ssDNA wrapping sequence programs protein corona composition. (**A**) Corona preparation. DNA-SWCNTs were incubated in human plasma, washed four times by centrifugation and PBS resuspension, and the bound corona was eluted with HFIP and analyzed by LC-MS/MS. (**B**) Overlap of enriched proteins across the three coronas, defined as log_2_FC > 0.585 against full plasma. (**C**) Mean log_2_FC of functionally defined protein modules. Error bars are s.e.m. The dotted line denotes no change against plasma. (**D**) Member-level log_2_FC for the coagulation modules: contact-activation proteins and fibrinogen complexes. Red: (AT)_15_-SWCNT. Black/gray: (CT)_15_-SWCNT. Blue: (GT)_15_-SWCNT.

We then compared the coronas at the level of functionally defined protein modules (**Figure 3C**). Forty-five proteins were enriched above 1.5-fold in all three coronas, constituting a wrapping-independent interaction core (Figure S9), including matrix Gla protein, insulin-like growth factor binding protein 3, versican, vitronectin, and selenoprotein P. The corona from (CT)_15_-SWCNTs carried the largest exclusive set of 26 proteins, dominated by lipoproteins, cytoskeletal/glycolytic proteins, coagulation factors, and the fibrinogen complex (Figure S10). While lipoprotein/lipid-transport proteins are enriched in all DNA-SWCNT coronas, (CT)_15_ wrapping enriched the lipoproteins the most broadly and strongly. Complement engagement was pathway specific. Lectin-pathway pattern-recognition proteins were enriched on (CT)_15_ comparably to or above other DNA wrapping, whereas classical-pathway components and several terminal/regulatory complement proteins were comparatively depleted. The (GT)_15_-SWCNT dominated block was enriched for classical-complement recognition, regulatory proteins and the matrix scaffold fibronectin (Figure S11). The (AT)_15_-SWCNT dominated block was the smallest, comprising contact-activation, neutrophil-derived protease cathepsin G, the NET-associated linker histones, and moderately enriched complement-associated proteins (Figure S12). Within the broader set of coagulation-associated proteins, while they are broadly represented in all three coronas, a functional split tracks intracellular retention (**Figure 3D**). Contact-activation proteins are consistently enriched in both high-retention, colloidally stable (AT)_15_– and (GT)_15_-SWCNTs, while fibrinogen complex proteins are most enriched in less stable, faster-clearing (CT)_15_-SWCNTs. This pattern maps onto the known sequence of competitive plasma protein adsorption, in which fibrinogen adsorbs early to anionic surfaces and is subsequently displaced by kininogen and other contact-system proteins.^52,53^ The (AT)_15_– and (GT)_15_ coronas carry the displacing proteins, e.g., histidine-rich glycoprotein and kininogen-1, and deplete fibrinogen. The (CT)_15_ corona has the opposite composition and thus does not support exchange from fibrinogen with contact activation proteins. Histidine-rich glycoprotein and kininogen-1 act as dysopsonins on nanoparticle surfaces and prolong particle residence,^54^ matching the high intracellular retention of (AT)_15_– and (GT)_15_-SWCNTs. Fibrinogen carries multiple binding domains and can bridge adjacent particles,^55^ leading to the substantial bundling of (CT)_15_-SWCNTs already evident at 1 h (**Figure 2f**).

### Wrapping Sequence Drives Divergent Proteomic Disruption and Lysosomal Engagement

Because complement, lipoprotein, and contact-activation proteins are recognized by different macrophage receptor families, the compositional divergence of the nanotube coronas suggests that each construct presents a distinct interfacial identity to the cell. To determine whether this divergence produces measurable differences in how macrophages handle each construct, we conducted time-resolved intracellular proteomics of macrophage lysates at 1, 6, and 24 h (**Figure 4A**, Figure S13).

**Figure 4.**
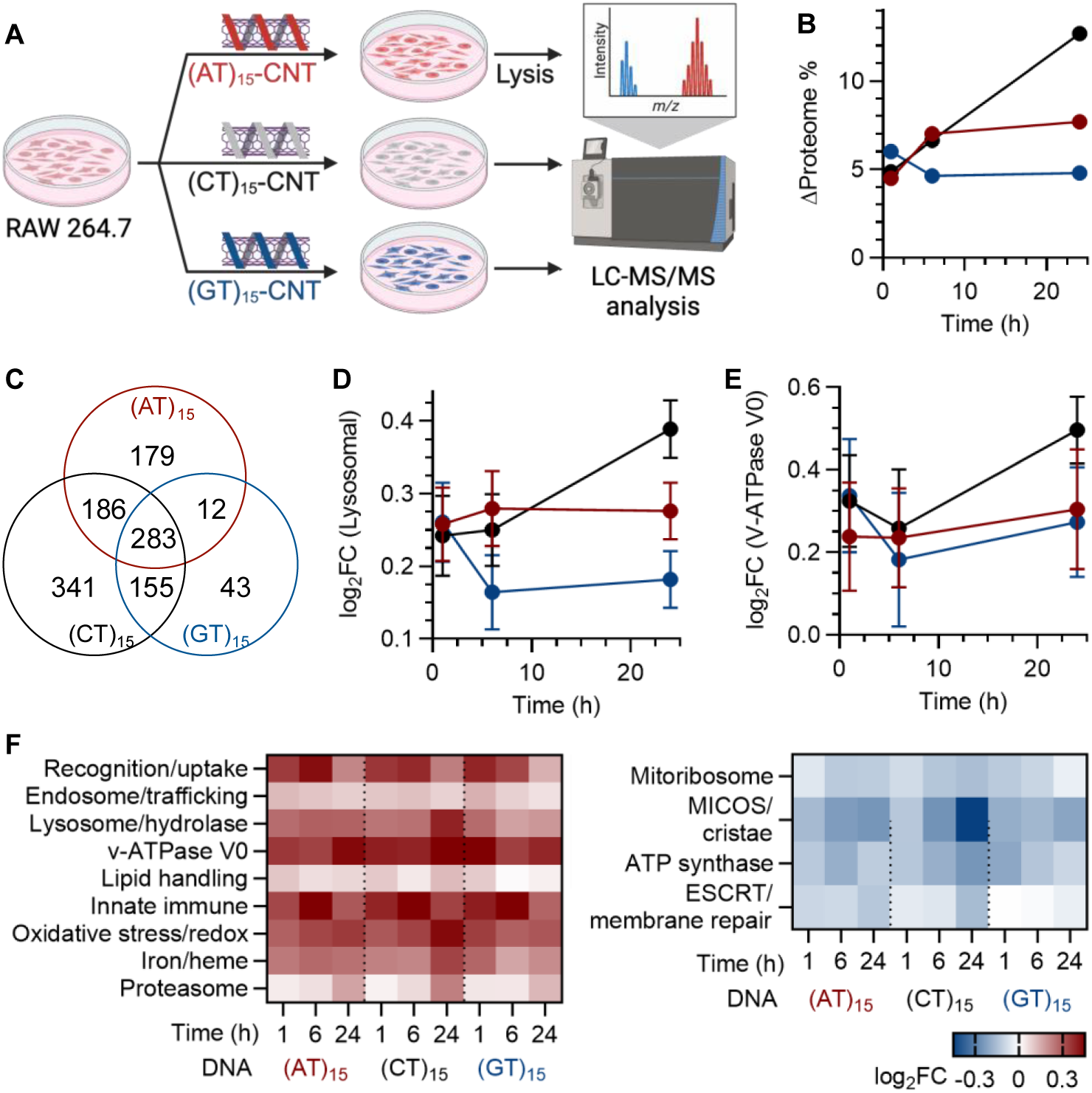
Wrapping sequence sets the extent of the macrophage proteome response. (**A**) Experimental design. RAW 264.7 macrophages were exposed to DNA-SWCNTs at 1 mgL^-1^ for 30 min, washed, and lysed at 1, 6, and 24 h for LC-MS/MS against a time-matched untreated control. (**B**) Overlap of enriched proteins across the three cell lysates, defined as log_2_FC > 0.585 against CNT-free control (**C**) Percentage of the quantifiable lysate proteome significantly altered at each time point. (**D**) Mean log_2_FC of a 16-member panel of lysosomal hydrolases and membrane proteins. (**E**) Mean log_2_FC of the v-ATPase V0 membrane sector. Red: (AT)_15_-SWCNT. Black/gray: (CT)_15_-SWCNT. Blue: (GT)_15_-SWCNT. (**F**) Mean log_2_FC of functionally defined protein panels across all nine conditions, ordered from most increased to most decreased. Panel membership and per-protein values are given in Table S5.

In the macrophage recognition machinery, receptors associated with lipoprotein and scavenger uptake, including CD36, LRP1, CD68, and COLEC12, were significantly elevated in all three constructs at comparable magnitude. Clathrin, AP2, and phagocytic actin components showed the same pattern, as did lipid handling downstream of uptake (Figure S14). Lysosomal acid lipase LIPA, the sterol transporters NPC1 and NPC2, and the acyl transferase SOAT1 rose in every construct, while de novo lipogenesis was suppressed (Figure S15). The construct carrying the most lipoprotein-rich corona therefore did not produce a distinct lipid-handling response. Complement and opsonin receptors showed their strongest response in (GT)_15_-SWCNT at 1 h, which matches the classical-complement composition of the (GT)_15_ corona.

We next interrogated how each DNA-SWCNT engages the endolysosomal degradation machinery, and whether its timing and severity track the optical and structural fate observed in **Figure 2**. Certain fractions of the quantifiable lysate proteome were significantly altered relative to time-matched control (**Figure 4B**). (AT)_15_-SWCNTs rose from 4.5% at 1 h to 7.0% at 6 h and plateaued at 7.7% by 24 h. (CT)_15_-SWCNTs began at 4.9% and increased to 12.7% by 24 h, the largest disruption of the three. (GT)_15_-SWCNTs peaked earliest at 6.0% at 1 h and resolved to 4.8% by 24 h. Considered cumulatively, (CT)_15_-SWCNTs altered 16.6% of the proteome over time, while the (AT)_15_ and (GT)_15_-SWCNTs reached comparable totals near 10% on different timing. Across the time course, 1,199 proteins were altered in at least one construct (**Figure 4C**). The (CT)_15_-specific set of 341 proteins exceeded the 283 altered in all three constructs, while only 12 proteins were altered in both (AT)_15_ and (GT)_15_ without also being altered in (CT)_15_.

These differences reflect engagement of the degradative compartment. A panel of 16 lysosomal hydrolases and membrane proteins was enriched among upregulated proteins, significantly in four of nine conditions and directionally consistent in all nine (**Figure 4D**). The mean panel response increased comparably across the three constructs at 1 h despite differences in internalized DNA-SWCNTs. By 24 h, the three had diverged. (AT)_15_ plateaued at +0.276, (GT)_15_ resolved to +0.182, and (CT)_15_ increased to +0.389, with all 16 panel members significant. Prosaposin, a lysosomal lipid-transfer protein, was the most strongly induced and was significant in all nine conditions. Lysosomal engagement thus diverges after uptake and follows the same ordering as intracellular nanotube loss quantified by the Raman G-band. Subunits of the v-ATPase V0 membrane sector were elevated in all nine conditions and were highest in (CT)_15_ at 24 h, which independently supports expansion of the endolysosomal compartment (**Figure 4E**). Lysosomal membrane integrity was maintained over 24 h, evidenced by unchanged or decreased levels of Galectin-3 and galectin-8, which accumulate on ruptured endolysosomal membranes, and depletion of the ESCRT membrane repair proteins (**Figure 4F**, Figure S16).

Beyond this divergence, the 283 proteins altered in all three constructs constitute a wrapping-independent core. The core included endosomal sorting machinery. TOM1L1, which recognizes ubiquitinated cargo on the endosomal membrane and directs it toward lysosomal degradation, was elevated in all nine conditions. Its paralogue TOM1 was elevated in all three constructs at 1 h and 6 h, and returned toward baseline by 24 h in (CT)_15_ and (GT)_15_. The late-endosomal marker CD63 was elevated in eight of nine conditions and reached its largest value in (CT)_15_ at 24 h. Sorting and priming machinery thus engaged early and resolved, whereas lysosomal capacity persisted. A second shared layer comprised the autophagy adaptor SQSTM1/p62 and the innate-immune proteins TLR2, NLRP3, and NFKB1, all of which peaked at 6 h and returned toward baseline by 24 h. Pathway enrichment identified ROS detoxification as the most consistent enrichment across all three sequences and at every time point (Figure S17).

### Oxidative and Proteostatic Output Follows the Degree of Lysosomal Engagement

We determined whether proteostatic and oxidative output followed the extent of lysosomal engagement (**Figure 4F**). Cytosolic proteolytic capacity followed it directly. Proteasome subunits were induced at 24 h in all three constructs, and the response was largest in (CT)_15_, followed by (AT)_15_ and (GT)_15_ (Figure S18). Mitochondrial proteins constitute a further wrapping-independent component of the response, and the response is one of depletion (Figure S19). Of 671 mitochondria-annotated proteins quantified, 111 were significantly altered in at least one construct and 28 in all three. Mitoribosomal subunits were significantly depleted in seven of nine conditions, and the MICOS and cristae scaffold proteins IMMT, PHB2, and CHCHD3 decreased in all nine. ATP synthase subunits also decreased. Respiratory-chain subunits were largely unchanged, with the exception of NDUFA8, which increased in all nine conditions. These changes indicate that mitochondrial translation and inner-membrane organization are engaged as part of the shared core response, and that their magnitude follows the same ordering as lysosomal engagement.

Functional assays resolved whether these proteomic differences correspond to measurable changes in oxidative and metabolic output, and the functional signature tracks the wrapping-dependent intracellular fate. (CT)_15_-SWCNT treated macrophages produced the highest general ROS (7-fold over control) and superoxide (3.5-fold), which matches the severe lysosomal engagement and the coordinated antioxidant and iron-handling response it provokes (**Figures 5A-B**). (AT)_15_-SWCNT treated cells showed moderate ROS and superoxide elevation. (GT)_15_-SWCNTs, in contrast, produced minimal general ROS, the lowest but significant superoxide rise (1.5-fold), and a significant (10-fold) rise in intracellular ATP over control (**Figure 5C**). The rise in intracellular ATP without a corresponding increase in general ROS points to a metabolic shift rather than a general oxidative one. The lysate proteome does not explain this phenotype through respiratory-chain abundance, since ATP synthase subunits decreased in (GT)_15_ and complex I subunits were unchanged. The ATP rise therefore reflects a change in metabolic flux, most plausibly towards glycolytic ATP production of the kind described in TLR-activated macrophages, this is inferred as flux was not measured directly. Antioxidant defenses were induced in all three constructs and at every time point. SOD1, the glutathione peroxidases GPX1 and GPX4, and thioredoxin reductase TXNRD1 were significantly elevated across the time course. SOD1, GPX1, and TXNRD1 reached their largest values in (CT)_15_ at 24 h. Antioxidant induction is therefore part of the shared core response and follows the same ordering as lysosomal engagement (Figure S17). Iron-handling proteins followed the same pattern. Ferritin light chain and peroxiredoxin 6 were elevated across the time course and reached their largest values in (CT)_15_ at 24 h (Figure S20).

**Figure 5.**
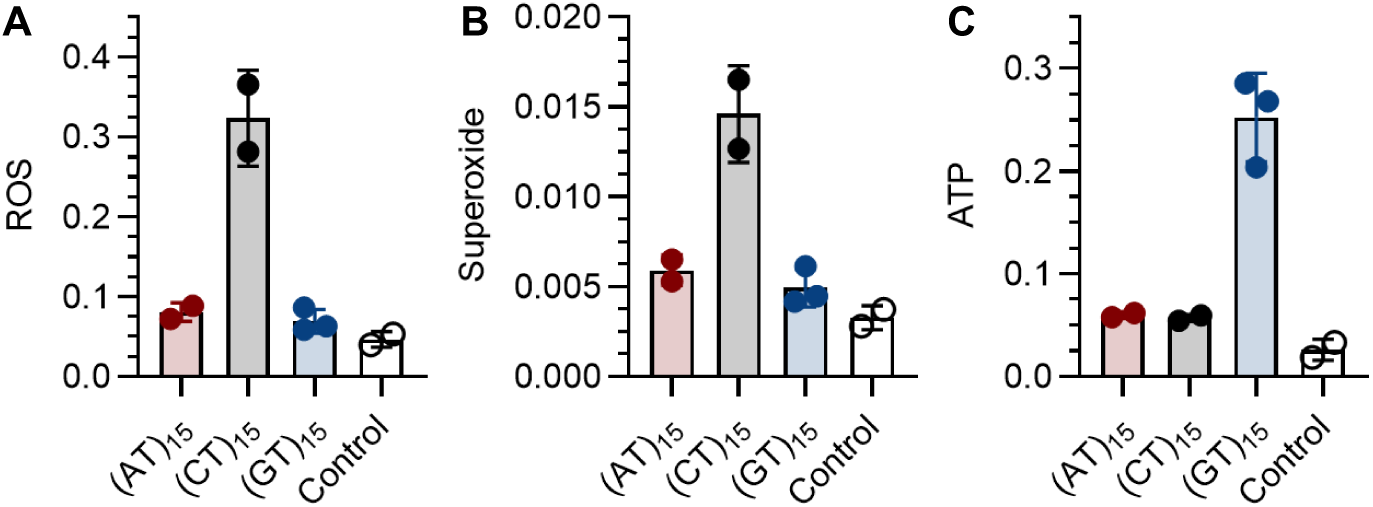
Functional oxidative and metabolic output tracks the wrapping sequence. RAW 264.7 macrophages were exposed to DNA-SWCNTs at 1 mg L^-1^ for 30 min, washed, and assayed at 24 h. (**A**) Total reactive oxygen species and (**B**) superoxide, measured with the ROS/superoxide detection assay and normalized to cell number. (**C**) Intracellular ATP, measured by luciferase assay and normalized to cell density. Bars are the mean of biological replicates, and error bars are std.d. Untreated cells served as the control.

## DISCUSSION

We demonstrate that the nucleotide-level surface identity of a DNA-wrapped SWCNT sets its intracellular fate. (AT)_15_-SWCNTs were internalized the most yet perturbed the proteome moderately and plateaued by 24 h. (CT)_15_-SWCNTs were internalized the least yet perturbed the proteome the most, and their disruption escalated through 24 h. (GT)_15_-SWCNTs perturbed the proteome earliest and resolved by 24 h to the smallest extent. Uptake magnitude therefore does not predict intracellular consequence. Nanotube integrity was most preserved by (AT)_15_ wrapping, but the least internalized sequence, (CT)_15_, provoked the largest rise in the defect density through time (135% rise in the Raman D-to-G ratio) and (GT)_15_ saw an initial increase in the defect ratio which did not change substantially through time. (AT)_15_– and (GT)_15_-SWCNTs exhibited comparable intracellular retention (71% and 79% of Raman G-band signals retained at 24 h, respectively), but the (CT)_15_-SWCNT signal was lost by over 60%. (CT)_15_-SWCNTs exhibit a large fluorescence peak width and broad distribution at 1 h, indicating substantial bundling from the earliest time point, whereas (AT)_15_– and (GT)_15_-SWCNTs remain comparatively individualized. (CT)_15_ is also the only construct in which lysosomal engagement increases through 24 h and the only construct that accumulates sidewall damage progressively through 24 hours. The compartment that (CT)_15_-SWCNTs occupy is both the most acidified by V0 sector abundance and the most oxidative, by ferritin and peroxiredoxin induction. The intracellular consequence is therefore set not by how much nanomaterial enters the cell but by how each surface wrapping and corona is recognized once inside. Aggregation state on arrival appears to determine how efficiently macrophages process the material.

Beneath the sequence-tuned differences lies a wrapping-independent response. It comprises particle engagement and vesicular sorting, represented by TOM1, TOM1L1, prosaposin, and mitochondrial proteostasis, represented by depletion of mitoribosomal and inner-membrane proteins. A second shared layer comprises TLR2, NLRP3, and SQSTM1/p62, which rise and resolve at a common 6 h maximum. The sequence effect is how far that response extends. These proteostatic responses parallel both the oxidative burden and the eventual structural fate of the SWCNTs. Severe structural damage to (CT)_15_-SWCNTs coincided with the highest cellular oxidative burden (7-fold ROS, 3.5-fold superoxide) and the most extensive proteomic disruption. In contrast, neither (AT)_15_– nor (GT)_15_-SWCNTs produced a significant rise in ROS or superoxide, and (GT)_15_ instead raised intracellular ATP, a metabolic rather than oxidative response. Consistent with this, neither construct showed significant structural damage temporally. A single ordering, (CT)_15_ > (AT)_15_ > (GT)_15_, describes proteome disruption, lysosomal engagement, proteasome induction, iron handling, mitochondrial depletion, and nanotube loss at 24 h. Uptake magnitude at 1 h follows the reverse ordering, and ATP rise in (GT)_15_-SWCNTs fits neither.

These findings refine how one should interpret corona composition as a predictor of cellular fate. The (CT)_15_ corona is the most compositionally distinct of the three, and (CT)_15_-SWCNTs are the outlier in every measurement we made. They accumulate the most sidewall damage, retain the least material, produce the highest oxidative output, and disrupt the proteome most. The (AT)_15_ and (GT)_15_ coronas are the most similar pair, and their intracellular responses differ in trajectory rather than in final extent. Across these three constructs, the degree of corona compositional divergence tracked the magnitude of the intracellular response, while the timing tracked the colloidal state of the material on arrival. The compositional feature that most distinguishes the (CT)_15_ corona is the lipoprotein and lectin-pathway recognition protein set. Because the coronas were formed in the same 10% human plasma used for cellular exposure, the corona profiled here is the one presented to the cell. The receptors, however, are murine, so the assignments remain inferred. Receptors for lipoprotein and scavenger uptake were elevated at comparable magnitude in all three constructs, which indicates that the shared recognition module we measure does not independently resolve the corona-specific engagement. The lysate proteome also averages over compartments, which is why we could not resolve the (GT)_15_ metabolic phenotype. We therefore treat the corona assignments as compositionally grounded hypotheses.

In summary, we found that the nucleotide-level ssDNA wrapping sequence is a determinant of intracellular fate, decoupled from nanotube uptake and bulk physicochemical properties, manifesting in construct-specific proteomic and oxidative dynamics. We propose that this acts through corona-mediated recognition, modulated by the colloidal state each construct adopts on arrival, since aggregation alters both the surface available for protein binding and the effective particle the cell encounters; sequence-resolved receptor engagement could not be directly observed here This positions the wrapping sequence as a practical design handle for tuning the intracellular behavior of carbon-based nanoprobes at fixed chirality, surface charge, and defect density parameters that current nanotoxicity frameworks rely on.^8^

## MATERIALS AND METHODS

### Materials

CoMoCAT SG65i raw SWCNT powder was purchased from Sigma-Aldrich. Sodium deoxycholate (SDC), sodium dodecyl sulfate (SDS), polyethylene glycol (PEG, 6 kDa), dextran (70 kDa), 2-propanol, and methanol were obtained from Sigma-Aldrich. ssDNA oligonucleotides (AT)_15_, (GT)_15_, and (CT)_15_ were purchased from Integrated DNA Technologies (IDT) at HPLC-purification grade. RAW 264.7 TIB-71 murine macrophages were obtained from ATCC (Manassas, VA). DMEM high-glucose, heat-inactivated FBS, HEPES, L-glutamine, penicillin/streptomycin, and amphotericin B were from Gibco. Cell Titer-Glo 2.0 (ATP assay) was from Promega. ROS/Superoxide Detection Assay kit (ab139476) was from Abcam. Bradford assay reagents were from Bio-Rad. Paraformaldehyde (4%) was from Thermo Fisher Scientific.

### SWCNT Dispersion Preparation and Aqueous Two-Phase Extraction

(6,5) chirality was enriched using a surfactant based aqueous two-phase extraction protocol^43^. Stock 1 is prepared by combining 42.3 g of 6 kDa polyethylene glycol (Thermo Scientific, Cat: A17541.0B, Lot. 10253547), 14.7 g of 70 kDa dextran (TCI, Cat: 9005-54-0, Lot. 7ALSL-MS), 1.47 g of sodium cholate (Sigma-Aldrich, Lot: 102640683), 3.351 g of sodium dodecyl sulfate (Sigma-Aldrich, Lot: 0000286470, ≥98.5%, 0.426 g of sodium chloride (Sigma-Aldrich, Lot: 0000275233, ≥99.0%), and 300 g of nanopore water. Stock 1 is mixed overnight at room temperature to ensure proper dissolution. The following day, 800 µL of CNT dispersion was combined with 1.581 mL of water and 15.18 mL of Stock 1. Mixture was thoroughly vortexed and centrifuged at 6000*g* for 5 min in a fixed-angle centrifuge (Eppendorf 5430R). The resulting top phase and interface were aspirated off, and the bottom phase was diluted 1:20 in 1 wt% sodium deoxycholate (aq). Purification was performed using dialysis membrane centrifugal filters (Amicon Ultra, Regenerated Cellulose, 100K NMWL, 15 mL, Lot: 0000433432). 15 mL of the diluted bottom phase was loaded onto the centrifugal filter and filtered at 2000*g* for 5 min. The process was repeated until a volume of 1 to 1.5 mL remained, thoroughly resuspending filter-bound nanotubes after each 5 min pulse. This filtration process was repeated for at least 3 cycles to reduce the polymer concentration by a factor of 10^3^. After 3 cycles, a small amount CNT suspension was removed and combined with methanol (≥99.9%, Sigma-Aldrich, Lot: 1003796521) in 1 to 2 volumetric ratio. The mixture was briefly vortexed and examined for turbidity resulting from residual polymer contamination. If turbidity was present, the purification cycle was repeated, and the sample was tested with methanol again. Once the CNT/methanol mixture was no longer turbid, the CNT suspension was examined using UV-vis-NIR absorption spectroscopy to quantify (6,5) concentration. After purification and concentration, functionalized CNT stock solutions were stored in the dark at room temperature.

### Surfactant-to-ssDNA Surface Exchange

Single-stranded DNA from Integrated DNA Technologies, DSPE-PEG 2000 (Avanti, Lot: 880128P-50MG-A-077), DMG-PEG 2000 (Avanti, Lot: 880151P-5G-A-027) were used as received. Polymer stock solutions were prepared as follows. For ssDNA, DNA was combined with DI water and pipetted thoroughly to form 10 mg/mL solutions. For modified PEG, the dry polymer powder was combined with water and bath-sonicated for 30 min (37 kHz, 60% power) to form a 5 mg/mL solution. To exchange the surfactant wrapping on the nanotube surface with the desired polymer wrapping agent, the following protocol was used^44^. In a 2 mL Eppendorf tube, 250 µL of the selected CNT stock was combined with 50 µL of either ssDNA or PEG stock solution. While vortexing at low speed, 1 mL of methanol was added slowly to precipitate nanotubes, followed by 50 uL of 5 M NaCl. Precipitated nanotubes were then removed from vortexing and centrifuged at 21,300*g* for 5 min. After centrifugation, the supernatant was aspirated and 250 µL of DI water was added. The CNT precipitate was then vigorously bath sonicated and pipetted until no visible particulate was visible. The cycle of polymer addition, precipitation, and resuspension in water was then repeated once more. After the second rewrapping, the polymer-wrapped nanotube suspension was then combined with 100 µL of additional polymer stock and diluted with 650 µL of DI water to form a 1 mL suspension of 1.0 mg/mL DNA or 0.5 mg/mL PEG, respectively. The resulting 1 mL suspensions were then tip sonicated at 30% power for 30 min. After tip sonication, polymer-wrapped CNT suspensions were loaded into a floating dialysis membrane (Float-A-Lyzer G2, MWCO: 1000 kDa, 1 mL) and placed in 1 L of nanopure water. Dialysis was performed overnight with bath exchange at 1 hr. and 4 hr. at room temperature. After dialysis, suspensions were characterized by UV-vis-NIR absorption, fluorescence, and zeta potential. Dialyzed nanotube suspensions were used in plasma incubation the same day as dialysis completion.

### Spectroscopic Characterization

For the absorption and fluorescence measurements, the nanotubes were dispersed in an aqueous solution of polymers or sodium deoxycholate as described in the earlier paragraph. All measurements were taken in Eppendorf UVette Cuvettes (Cat: 952010051). Absorbance spectra of nanotubes were collected by a V-780 UV-Visible-NIR spectrophotometer (Jasco, Inc.) with the following parameters: UV-Vis bandwidth of 2.0 nm, NIR bandwidth of 4.0 nm, range of 300–1,350 nm, scan speed of 1,000 nm/min. The extinction coefficient A_910_ = 0.02554 L mg^−1^ cm^−1^ was used to determine SWCNT concentration. Dispersion yield was calculated as the mass fraction of SWCNTs recovered in the ssDNA-SWCNT supernatant relative to the starting DOC-SWCNT stock. The optical density at (6,5) E_11_ was adjusted to 0.15 to avoid inner filter effects in the fluorescence measurements. Fluorescence emission spectra of nanotubes were acquired by NS Super (Applied Nano Fluorescence, Houston TX, USA). The samples were excited at 575 nm (bandwidth of 10 nm, 10-20 mW, averaging: 3, integration time: 2 seconds). Following the fluorescence spectral acquisition, the data were processed using custom MATLAB code that applied the spectral corrections and background subtraction, and the fluorescence emission peaks were fitted with Lorentzian functions.

### Zeta potential

An aqueous solution of DNA-SWCNTs in nanopure water with an optical density at (6,5) E_11_ of 0.15 was transferred into a DTS1070 folded capillary Zetasizer Cell (Malvern, M00049654). Zeta potential values were acquired with the Malvern Zetasizer Nano Z. Instrument parameters for our dispersant are as follows: 25°C, 0.8872 cP viscosity, refractive index of 1.330, and 78.5 dielectric constant. Equilibrium time is set to 120 seconds. The zeta potentials were calculated with the Smoluchowski model.

### Confocal Raman microscopy

Solid state nanotube samples were prepared from aqueous dispersions prior to Raman characterization. Briefly, 100 µL of nanotube suspension (optical density of 1 at (6,5) E_11_) dispersed in sodium deoxycholate (1 wt%, aq) was combined with 1,900 µL of ethanol to destabilize the surfactant shell and promote nanotube aggregation. The mixture was centrifuged at 13,000 *g* for 5 minutes, after which the supernatant was discarded. The remaining pellet was deposited onto clean glass substrates and dried overnight in a fume hood at room temperature. Raman spectra of the dried SWCNT film were acquired on a QONTOR Confocal Raman spectrometer (Renishaw) under 785 nm excitation laser. Spectra were collected using a 1,200 lines mm^-1^ diffraction grating, an incident laser power of 40 mW, a 1s integration time, and 25 accumulations per spectrum, with the excitation laser beam focused onto the sample through a 50x objective. Raman spectra of fixed cells were acquired using a Renishaw inVia confocal Raman microscope equipped with a 50× long working distance objective (LM Plan 50/0.6), 785 nm laser (30 mW at the sample stage), and 1200 lines/mm grating. Individual cells were raster-scanned at 1×1 μm^2^ intervals with 2 s integration. Spectra were recorded over the 830– 1,900 cm^-1^ spectral acquisition range, measuring both the D band (∼1,300 cm^-1^) and the G band (∼1,585 cm^-1^) of DNA-SWCNTs. Multiple spatial positions across each sample were measured to account for any film heterogeneity. Background subtraction was performed in WiRE 5.2 software (Renishaw) using the standard baseline correction modality. The D/G intensity ratio was calculated from the baseline corrected spectra to quantify relative defect densities of the nanotube samples.

### Cell Culture

RAW 264.7 murine macrophages (ATCC TIB-71) were maintained at 37°C, 5% CO_2_ in D10 medium: high-glucose DMEM supplemented with 10% heat-inactivated FBS, 2.5% HEPES, 1% L-glutamine, 1% penicillin/streptomycin, and 0.2% amphotericin B (all Gibco).

### SWCNT Treatment to Cells

Cells were seeded at 5.26 × 10^4^ cells cm^−2^ on 35 mm glass-bottom microwell dishes (MatTek) for microscopy or in 96-well plates for functional assays and cultured overnight prior to SWCNT treatment. For SWCNT exposure, culture medium was replaced with 1 mg L^−1^ ssDNA-SWCNT diluted in 10% Human Plasma and cells were incubated for 30 min. SWCNT-containing medium was then removed, cells were washed twice with sterile PBS (Gibco), and fresh D10 culture medium was added (Control cell samples were incubated with a blank SWCNT solution which comprised of 10% Human Serum but no SWCNTs). Time points are defined relative to this wash step. For Raman microscopy, cells were fixed with 4% paraformaldehyde in PBS (15 min, room temperature), washed three times with PBS, and maintained in PBS during imaging.

### Hyperspectral NIR Fluorescence Imaging

Hyperspectral NIR fluorescence image stacks were acquired using the IMA system (Photon Etc.) coupled to an Olympus IX-73 inverted microscope with a 20× IR objective (LCPlan N, 20×/0.45 IR; Olympus). A 577 nm laser provided excitation. Live cell imaging was performed on a stage-top incubator (Oko Lab) that maintained 37°C and 5% CO_2_ throughout the spectral acquisition. Hyperspectral cubes were processed in MATLAB to extract (6,5) peak emission intensity, center wavelength, and FWHM on a pixel-by-pixel basis. For long-term tracking (1–72 h), ROI-level spectral metrics were extracted from individual cells.

### ATP quantification

Intracellular ATP was measured in 96-well plates using the Cell Titer-Glo 2.0 assay (Promega) following manufacturer instructions. Luminescence was recorded on a BioTek Synergy H4 microplate reader 10 min after reagent addition. ATP concentrations were calculated against a known ATP standard curve and normalized to cell density.

### ROS and superoxide

Reactive oxygen species and superoxide levels were measured using the Abcam ROS/Superoxide Detection Assay (ab139476) per manufacturer protocol. Fluorescence was measured at Ex/Em 520/605 nm using a BioTek Synergy H4 microplate reader and reported as normalized intensity relative to cell number.

### Protein quantification

The Detergent Compatible Bradford Assay Reagent (Thermo Scientific, Lot: ZE390357) was warmed to room temperature before use. An albumin standard of 2.0 mg/mL in 0.9% NaCl (Thermo Scientific, Lot: ZE392797) was used to prepare a range of standard concentration samples as instructed. The standard and the protein samples were plated in volumes of 150 µL in a flat-bottom Corning 96-well plate. 150 µL of the Bradford reagent was dispensed rapidly into each well. After 5 minutes of incubation at room temperature and in the dark, the absorbance at 595 nm was read using a Biotek Synergy HT Multi-Mode Microplate Reader.

### Viability

Cell viability was assessed using the Tali image-based cytometry kit (Thermo Fisher) on an ECHO R2-Resolve fluorescence microscope. Cells were stained per protocol and analyzed for live/dead ratios in triplicate.

### Plasma Incubation and Protein Corona Extraction

Pooled human plasma (Blood derived, K2 EDTA, Innovative Research, Inc.) was thawed and centrifuged at 13,000*g* at 4 °C for 10 min. After centrifugation, the resulting fatty surface layer was aspirated, and 600 uL of plasma supernatant was combined with 300 µL of 10x PBS, polymer-wrapped CNT stock, and water to form a 3 mL mixture of (6,5) CNT solution with the optical density at (6,5) E_11_ of 0.15. The resulting (6,5) nanotube concentration is approximately 2 µg mL^-1^. The resulting mixture was placed on a Fisherbrand Multi-Platform Shaker in a closed container at 200 speed for 120 min. The incubated mixture was divided into 3 x 1 mL aliquots in Eppendorf Protein LoBind Tubes (1.5 mL, Lot: O221182K) and centrifuged at 21,300 *g* at 4 °C for 25 min. After centrifugation, a visible CNT pellet appears at the bottom of the container. The supernatant is gently aspirated off, and pellets are recombined by transferring 1250 µL 1X PBS between the 3 aliquot tubes to collect the pellets into a single Eppendorf tube. This process is repeated two more times. For final resuspension, pellets are suspended in 200 µL of 1,1,1,3,3,3-hexafluoro-2-propanol (HFIP) (>99%, Sigma-Aldrich, Lot: 1003743061), transferred to a PTFE membrane filter tube (Ultrafree-MC Centrifugal Filters, MilliporeSigma, 200 µm, Cat: UFC30LG25, Lot: N5BB27060), and centrifuged at 12,000*g* for 10 min. The HFIP elution step is repeated once more, resuspending the nanotube pellet in the supernatant chamber and centrifuging. Protein-containing HFIP elution is transferred to a fresh Protein LoBind tube, using additional HFIP to thoroughly remove all proteins from the filter tube. The protein suspension was then stored at –80 °C until digestion.

### Tryptic Digestion

All samples were dried via SpeedVac before using the Micro S-Trap (Protifi) columns for tryptic digestion as per the manufacturer’s instructions. Samples were resuspended in 23 µL of 1x sodium dodecyl sulfate Lysis buffer (5% sodium dodecyl sulfate, 50 mM TEAB) and shaken at 2000 rpm for 10 min to lyse the samples and dissolve proteins. Samples were reduced with TCEP to a final concentration of 5 mM and incubated at 55°C for 15 minutes. Reduced samples were then alkylated by the addition of Iodoacetamide (IAA) to a final concentration of 10 mM in solution and incubated at room temperature in the dark. Samples were acidified by the addition of 27.5% phosphoric acid to a final concentration of approximately 2.5% and vortexed to mix. 165 µL of binding buffer (100 mM TEAB (final) in 90% methanol) was added to each tube and shaken at 2000 rpm for 5 mins to mix. The contents of each tube were transferred to an individual S-Trap column placed in a 1.7 mL receiver tube. These tubes were spun at 4000 × g for 1 minute and the flow through was discarded. Trapped proteins were washed three times with wash buffer (100 mM TEAB (final) in 90% methanol), with the flow-through of each wash step discarded. MS grade trypsin protease (Thermo Scientific) was added to each trap column in a 1:50 trypsin to protein ratio dissolved in 20 µL of 50 mM TEAB. The S-Trap columns were incubated in a water bath at 37°C overnight. On the following day, digested peptides were eluted sequentially by 50 mM TEAB, 0.1% formic acid, and 50% acetonitrile. Pooled eluted peptides were dried via SpeedVac. Samples were resuspended in 25 µl of 5% acetonitrile, 0.1% formic acid, shaken at 2000 rpm for 10 minutes, centrifuged at 21 kg for 10 minutes, and the supernatant was collected for LC-MS data acquisition.

### Tandem Mass Tag (TMT) Label Tagging

A TMT 18-plex 0.5mg Mass Labeling Kit (Thermo Scientific) was used to isobarically label peptide samples per the manufacturer’s instructions. Immediately before using, the TMT label reagents were equilibrated to room temperature. 20 µL of anhydrous ACN was added to each tube to dissolve the reagent with occasional vortexing for 5 minutes. The protein digest was then resuspended in 10 µL of 100 mM TEAB. The contents of the TMT Reagent vial were transferred to the tubes containing the protein digest in solution. Samples were incubated at room temperature for 1 hour. To quench the reaction, 5 µL of 5% hydroxylamine was added to each sample and incubated at room temperature for 15 minutes. All labeled samples were combined, vortexed and divided into three microcentrifuge tubes for quantitative fold-change analysis.

### Reversed Phase-Reversed Phase Fractionation

A High pH Reversed-Phase Peptide Fractionation Kit (Thermo Scientific) was used to fractionate TMT-labeled peptides per the manufacturer’s instructions. The spin columns were placed into a collection tube and conditioned by centrifugation at 5,000 g for 2 minutes to remove the storage solution and pack the resin material. To wash the columns, 300 µL of anhydrous acetonitrile was added, and the spin column was centrifuged at 5,000 g for 2 minutes. This wash step was repeated with the flow-through of the wash discarded. The column was then washed twice, as in the previous step, with 0.1% TFA to complete the column conditioning. Labeled proteolytic digests were dissolved in 100 µL of 0.1% TFA and combined to a total of 300 µL. The conditioned spin column was placed into a new 2.0 mL receiver tube, and the 300 µL resuspended protein digest was loaded onto the column. The column was centrifuged at 3,000 g for 2 minutes, and the flow-through was retained. This step was repeated with water loaded onto the column, and the flow-through was collected as the “first wash” fraction. The “second wash” fraction was collected using the first elution solution to remove unreacted TMT reagent. The column was placed in a new receiving tube and 300 µL of the remaining gradient elution solutions were used with each fraction being collected by centrifugation at 3,000 g for 2 minutes. Each fraction was dried via SpeedVac, and the dried samples were resuspended in 25 µL of 0.1% formic acid, shaken at 2,000 rpm for 10 minutes, and centrifuged at 21 kg for 10 minutes before being placed in sample vials of 15 µL for LC-MS analysis.

### LC-MS/MS Data Acquisition

An externally calibrated Thermo Exploris 480 (high-resolution electrospray tandem mass spectrometer) was used in conjunction with Vanquish Neo nano LC System. 1 μg of each sample was aspirated into a 50 μL loop and loaded onto the trap column (Thermo µ-Precolumn 5 mm, with nanoViper tubing 30 µM i.d. × 10 cm). The flow rate was set to 300 nL/min for separation on the analytical column (EASY-Spray™ PepMap™ Neo UHPLC Column, 50 cm long, 75-micron internal diameter, C18, reverse phase). Mobile phase A was composed of 99.9% H2O (EMD Omni Solvent), and 0.1% formic acid and mobile phase B was composed of 80% ACN, and 0.1% formic acid. A 90-minute linear gradient from 5% to 55% B was performed. The LC eluent was directly nanosprayed into Exploris 480 mass spectrometer (Thermo Scientific). During the chromatographic separation, the Exploris 480 plus was operated in a data-dependent mode and under direct control of the Thermo Excalibur 4.7.69.37 Software (Thermo Scientific). MS data was acquired using the following parameters: data-dependent scan with a 3 second cycle time per full scan (400 to 1500 m/z) at 60,000 resolutions. MS2 was acquired at 15,000 resolutions. Ions with a single charge or charges of more than 8, as well as unassigned charges, were excluded. An auto-dynamic exclusion window was used. All measurements were performed at room temperature.

### LC-MS/MS Data Analysis

Resultant raw files were searched with Proteome Discoverer 3.2 using the Chimerys search engine with a *Homo sapiens* FASTA database and a *Mus musculus* FASTA for protein corona and cell lysates, respectively. A 20-ppm mass tolerance for parent ion and 0.02Da mass tolerance for the fragment ion were used. Analyses of individual proteins and of protein panels were restricted to proteins identified by at least 10 peptide-spectrum matches, since the pooled reference channel is measured once per TMT set and ratios for sparsely identified proteins carry correspondingly higher uncertainty. Proteome-wide fractions in Figure 4b were computed without this restriction.

Lysate samples were distributed across two TMT 18-plex sets. A pooled reference sample, comprising equal contributions from all samples in the experiment, was included as a bridge channel in both sets. Untreated control samples were included in the first set. All quantification was performed on the ratio of each sample channel to the pooled reference channel within its own set, which places all samples on a common scale across the two sets. For each protein, normalized abundances were converted to log2 ratios against the pooled reference channel within the same TMT set. Treated conditions were compared with untreated control by two-sample t-test, and P values were corrected across all quantified proteins by the Benjamini-Hochberg procedure. Proteins were called significantly altered at q < 0.05 and |log2 fold change| > 0.585.

### Statistical Analysis

All statistical analyses were performed in OriginPro 2024b. Data normality was assessed by the Shapiro-Wilk test, and variance homogeneity by Levene’s test. Where assumptions were met, significance was evaluated by two-sample t-test (pairwise comparisons) or one-way ANOVA with Tukey’s post hoc test (multiple comparisons). Where assumptions were violated, appropriate nonparametric tests were applied. A false discovery rate correction was applied for proteomic comparisons. Statistical significance thresholds: *p < 0.05, **p < 0.01, ***p < 0.001. All data represent the mean of at least n = 3 independent biological replicates unless otherwise noted. All data shown as mean ± s.d. unless otherwise specified.

## AUTHOR CONTRIBUTIONS

A.N. and M.K. conceived the study, performed data analysis and interpretation, and wrote the manuscript. A.N. performed SWCNT preparation, physicochemical characterization, cell culture, NIR hyperspectral imaging, and functional assays. J.M performed the proteomics experiments; T.X. and D.K. contributed to NIR and Raman imaging experiments. M.K. provided project supervision, resources, and funding. All authors reviewed and approved the final version.

## DATA AVAILABILITY

All raw data, analysis code, and proteomic datasets supporting the findings of this study will be deposited in a publicly accessible repository upon acceptance. Processed data and key analysis scripts are available from the corresponding author upon reasonable request.

## REFERENCES

1. Mitchell, M. J. et al. Engineering precision nanoparticles for drug delivery. Nat. Rev. Drug Discov. 20, 101–124 (2021).

2. Hong, G., Antaris, A. L. & Dai, H. Near-infrared fluorophores for biomedical imaging. *Nat*. Biomed. Eng. 1, 0010 (2017).

3. Welsher, K. et al. A route to brightly fluorescent carbon nanotubes for near-infrared imaging in mice. Nat. Nanotechnol. 4, 773–780 (2009).

4. Kruss, S. et al. Carbon nanotubes as optical biomedical sensors. Adv. Drug Deliv. Rev. 65, 1933– 1950 (2013).

5. Ackermann, J., Metternich, J. T., Herbertz, S. & Kruss, S. Biosensing with Fluorescent Carbon Nanotubes. Angew. Chem. Int. Ed. 61, e202112372 (2022).

6. Ryan, A. K., Israel, A., Stefoni, M. C., Ferraz, C. & Williams, R. M. Rational Design of Optical Single-Walled Carbon Nanotube-Based Nanosensors with Biological Recognition Elements. Adv. Sens. Res. 5, e00076 (2026).

7. Cohen, Z. & Williams, R. M. Single-Walled Carbon Nanotubes as Optical Transducers for Nanobiosensors In Vivo. ACS Nano 18, 35164–35181 (2024).

8. Kim, M. et al. Human and environmental safety of carbon nanotubes across their life cycle. Nat. Rev. Mater. 9, 63–81 (2023).

9. Nadeem, A. et al. Spectral Fingerprinting of Engineered Nanomaterials for Precision Biosensing. ACS Nano 20, 3921–3943 (2026).

10. Shvedova, A. A. et al. Unusual inflammatory and fibrogenic pulmonary responses to single-walled carbon nanotubes in mice. Am. J. Physiol.-Lung Cell. Mol. Physiol. 289, L698–L708 (2005).

11. Li, Z. et al. Cardiovascular Effects of Pulmonary Exposure to Single-Wall Carbon Nanotubes. Environ. Health Perspect. 115, 377–382 (2007).

12. Portilla, Y. et al. Different coatings on magnetic nanoparticles dictate their degradation kinetics in vivo for 15 months after intravenous administration in mice. J. Nanobiotechnology 20, 543 (2022).

13. Roxbury, D., Jagota, A. & Mittal, J. Sequence-Specific Self-Stitching Motif of Short Single-Stranded DNA on a Single-Walled Carbon Nanotube. J. Am. Chem. Soc. 133, 13545–13550 (2011).

14. Tu, X., Manohar, S., Jagota, A. & Zheng, M. DNA sequence motifs for structure-specific recognition and separation of carbon nanotubes. Nature 460, 250–253 (2009).

15. Salem, D. P. et al. Chirality dependent corona phase molecular recognition of DNA-wrapped carbon nanotubes. Carbon 97, 147–153 (2016).

16. Krug, H. F. Nanosafety Research—Are We on the Right Track? Angew. Chem. Int. Ed. 53, 12304–12319 (2014).

17. Nelissen, I. et al. Improving Quality in Nanoparticle-Induced Cytotoxicity Testing by a Tiered Inter-Laboratory Comparison Study. Nanomaterials 10, 1430 (2020).

18. Ong, K. J. et al. Widespread Nanoparticle-Assay Interference: Implications for Nanotoxicity Testing. PLoS ONE 9, e90650 (2014).

19. Nel, A. E. et al. Understanding biophysicochemical interactions at the nano–bio interface. Nat. Mater. 8, 543–557 (2009).

20. Huang-Zhu, C. A. & Van Lehn, R. C. Engineering the nano-bio interface: challenges and opportunities for predicting the surface properties of monolayer-protected nanoparticles. Mater. Horiz. 12, 6428–6439 (2025).

21. Cedervall, T. et al. Understanding the nanoparticle–protein corona using methods to quantify exchange rates and affinities of proteins for nanoparticles. Proc. Natl. Acad. Sci. 104, 2050–2055 (2007).

22. Salvati, A. et al. Transferrin-functionalized nanoparticles lose their targeting capabilities when a biomolecule corona adsorbs on the surface. Nat. Nanotechnol. 8, 137–143 (2013).

23. Ngo, W. et al. Identifying cell receptors for the nanoparticle protein corona using genome screens. Nat. Chem. Biol. 18, 1023–1031 (2022).

24. Lesniak, A. et al. Effects of the Presence or Absence of a Protein Corona on Silica Nanoparticle Uptake and Impact on Cells. ACS Nano 6, 5845–5857 (2012).

25. Tenzer, S. et al. Rapid formation of plasma protein corona critically affects nanoparticle pathophysiology. Nat. Nanotechnol. 8, 772–781 (2013).

26. Vu, V. P. et al. Immunoglobulin deposition on biomolecule corona determines complement opsonization efficiency of preclinical and clinical nanoparticles. Nat. Nanotechnol. 14, 260–268 (2019).

27. Xiao, W. et al. The protein corona hampers the transcytosis of transferrin-modified nanoparticles through blood–brain barrier and attenuates their targeting ability to brain tumor. Biomaterials 274, 120888 (2021).

28. Pinals, R. L., et al. Protein Corona Selectivity on Carbon Nanotube Biosensors is Driven by Surface Coating and Chirality. Preprint at 10.26434/chemrxiv.10001554/v1 (2026).

29. Miller, J., Coscia, A., Nadeem, A. & Kim, M. Structure-affinity correlations and separable optical activity in carbon nanotube protein coronas. Preprint at 10.64898/2026.07.16.739030 (2026).

30. Hwang, I.-J., Gray, J. S. & Kim, M. Chemically Engineered Carbon Nanotubes Map Class-Selective Metabolite Enrichment from Human Plasma. Preprint at 10.64898/2026.06.02.729405 (2026).

31. Walkey, C. D. et al. Protein Corona Fingerprinting Predicts the Cellular Interaction of Gold and Silver Nanoparticles. ACS Nano 8, 2439–2455 (2014).

32. Pinals, R. L., et al. Quantitative Protein Corona Composition and Dynamics on Carbon Nanotubes in Biological Environments. Angew. Chem. Int. Ed. 59, 23668–23677 (2020).

33. Mukherjee, S. P. et al. Macrophage sensing of single-walled carbon nanotubes via Toll-like receptors. Sci. Rep. 8, 1115 (2018).

34. Bussy, C. et al. Intracellular fate of carbon nanotubes inside murine macrophages: pH-dependent detachment of iron catalyst nanoparticles. Part. Fibre Toxicol. 10, 24 (2013).

35. González-García, L. E. et al. Nanoparticles Surface Chemistry Influence on Protein Corona Composition and Inflammatory Responses. Nanomaterials 12, 682 (2022).

36. Zheng, M. et al. DNA-assisted dispersion and separation of carbon nanotubes. Nat. Mater. 2, 338–342 (2003).

37. Zheng, M. et al. Structure-Based Carbon Nanotube Sorting by Sequence-Dependent DNA Assembly. Science 302, 1545–1548 (2003).

38. Nalige, S. S. et al. Fluorescence changes in carbon nanotube sensors correlate with THz absorption of hydration. Nat. Commun. 15, 6770 (2024).

39. Choi, J. H. & Strano, M. S. Solvatochromism in single-walled carbon nanotubes. Appl. Phys. Lett. 90, 223114 (2007).

40. Kruss, S. et al. Neurotransmitter Detection Using Corona Phase Molecular Recognition on Fluorescent Single-Walled Carbon Nanotube Sensors. J. Am. Chem. Soc. 136, 713–724 (2014).

41. Zhang, J. et al. Molecular recognition using corona phase complexes made of synthetic polymers adsorbed on carbon nanotubes. Nat. Nanotechnol. 8, 959–968 (2013).

42. Khripin, C. Y., Manohar, S., Zheng, M. & Jagota, A. Measurement of Electrostatic Properties of DNA-Carbon Nanotube Hybrids by Capillary Electrophoresis. J. Phys. Chem. C 113, 13616– 13621 (2009).

43. Subbaiyan, N. K. et al. Role of Surfactants and Salt in Aqueous Two-Phase Separation of Carbon Nanotubes toward Simple Chirality Isolation. ACS Nano 8, 1619–1628 (2014).

44. Streit, J. K., Fagan, J. A. & Zheng, M. A Low Energy Route to DNA-Wrapped Carbon Nanotubes via Replacement of Bile Salt Surfactants. Anal. Chem. 89, 10496–10503 (2017).

45. Nadeem, A. et al. Enhancing Intracellular Optical Performance and Stability of Engineered Nanomaterials via Aqueous Two-Phase Purification. Nano Lett. 23, 6588–6595 (2023).

46. Choi, J. H. & Strano, M. S. Solvatochromism in single-walled carbon nanotubes. Appl. Phys. Lett. 90, 223114 (2007).

47. Dresselhaus, M. S., Jorio, A. & Saito, R. Characterizing Graphene, Graphite, and Carbon Nanotubes by Raman Spectroscopy. Annu. Rev. Condens. Matter Phys. 1, 89–108 (2010).

48. Nadeem, A., Kindopp, A., Junge, E., Rahmani, M. & Roxbury, D. Enhancing optical properties and stability of DNA-functionalized carbon nanotubes with cryoprotectant-mediated lyophilization. Carbon 248, 121159 (2026).

49. Nadeem, A., Lyons, S., Kindopp, A., Jamieson, A. & Roxbury, D. Machine Learning-Assisted Near-Infrared Spectral Fingerprinting for Macrophage Phenotyping. ACS Nano 18, 22874–22887 (2024).

50. Gravely, M. et al. Aggregation Reduces Subcellular Localization and Cytotoxicity of Single-Walled Carbon Nanotubes. ACS Appl. Mater. Interfaces 14, 19168–19177 (2022).

51. Schöppler, F., Rühl, N. & Hertel, T. Photoluminescence microscopy and spectroscopy of individualized and aggregated single-wall carbon nanotubes. Chem. Phys. 413, 112–115 (2013).

52. Nienhaus, K. & Nienhaus, G. U. Mechanistic Understanding of Protein Corona Formation around Nanoparticles: Old Puzzles and New Insights. Small 19, 2301663 (2023).

53. Schmaier, A. H. et al. The effect of high molecular weight kininogen on surface-adsorbed fibrinogen. Thromb. Res. 33, 51–67 (1984).

54. Fedeli, C. et al. The functional dissection of the plasma corona of SiO_2_ –NPs spots histidine rich glycoprotein as a major player able to hamper nanoparticle capture by macrophages. Nanoscale 7, 17710–17728 (2015).

55. Emilsson, G. et al. The *In Vivo* Fate of Polycatecholamine Coated Nanoparticles Is Determined by a Fibrinogen Enriched Protein Corona. ACS Nano 17, 24725–24742 (2023).

