## Supporting Information for "DNA Sequence-Programmed Protein Coronas Determine Intracellular Fate and Proteostatic Stress of Carbon Nanotubes"

**Table S1**. UV-Vis-NIR spectroscopy results for pre- and post-aqueous two-phase extraction (ATPE) sorting processes.

| Metric | Pre-ATPE | Post-ATPE |
|---|---|---|
| (6,5) E11 Integrated Area | 57 % | 75% |
| (6,5) Resonance ratio at 991nm | 66% | 96% |
| Background OD fraction at (6,5) peak | 34% | 4% |

**Table S2.** Quantification of individual chiralities in the pre and post ATPE sorted samples.

| Chirality | λ (nm) | % in pre-ATPE | % in post-ATPE |
|---|---|---|---|
| (6,4) | 886 | 10 % | 9% |
| (6,5) | 991 | 57 % | 75% |
| (7,5) | 1050 | 15% | 2.4 % |
| (8,4) | 1129 | 9 % | 3 % |

**Table S3.** Quantification of (6,5) Center wavelength after the process of rewrapping showing significant red shifting of ssDNA wrappings from initial DOC dispersion.

| Non-covalent wrapping | (6,5) Center Wavelength (nm) |
|---|---|
| DOC | 985 |
| $(AT)_{15}$ | 992 |
| $(CT)_{15}$ | 996 |
| $(GT)_{15}$ | 998 |

**Table S4.** Intracellular Raman spectroscopy information showing how each wrapping type changes intracellular processing of these nanomaterials.

| Wrapping | G-band 1h (cts) | G-band 24h (cts) | % Retained | D/G 1h | D/G 24h | Δ D/G (%) |
|---|---|---|---|---|---|---|
| $(AT)_{15}$ | 926 | 656 | 70.8% | 0.039 | 0.035 | −10.3% |
| $(CT)_{15}$ | 446 | 176 | 39.5% | 0.129 | 0.303 | +134.9% |
| $(GT)_{15}$ | 530 | 420 | 79.2% | 0.235 | 0.258 | +9.8% |

**Table S5.** Cellular functional assay results for macrophages treated with DNA-SWCNTs.

| Assay | $(AT)_{15}$ | $(CT)_{15}$ | $(GT)_{15}$ | Control |
|---|---|---|---|---|
| ATP (lum./cell density) | 0.060 | 0.057 | 0.251** | 0.026 |
| ROS (norm. intensity) | 0.082 | 0.324*** | 0.071 | 0.045 |
| Superoxide (norm. intensity) | 0.005 | 0.014** | 0.005 | 0.004 |
| BCA Protein ($\times 10^{-4}$ abs.) | 0.01 | 0.27* | 0.55** | 0.02 |

**p < 0.01, ***p < 0.001, *p < 0.05

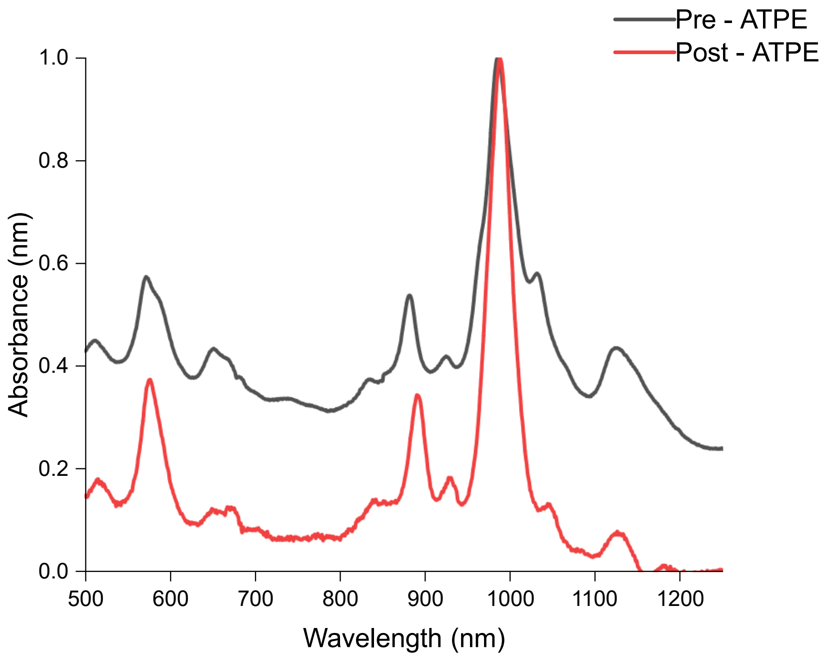

**Figure S1.** UV-vis-NIR absorption spectra of the CoMoCAT SG65i dispersion before and after ATPE. Enrichment raises the (6,5) $E_{11}$ integrated optical density from 57% to 75% and reduces the non-resonant background fraction at the (6,5) peak from 34% to 4%.

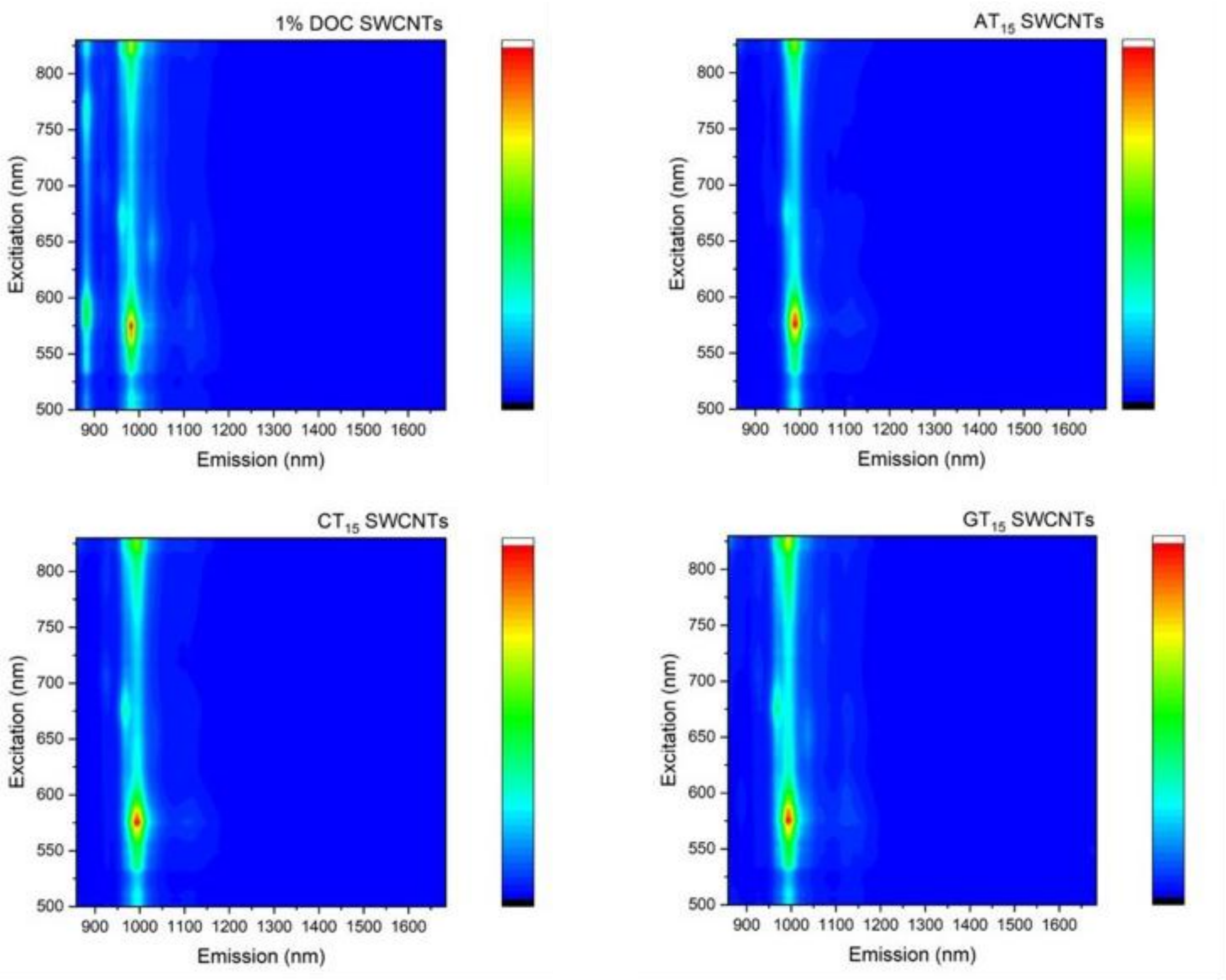

**Figure S2.** Two-dimensional excitation-emission fluorescence maps. Fluorescence maps of $(AT)_{15}$-, $(CT)_{15}$-, and $(GT)_{15}$-SWCNTs confirm (6,5) chirality enrichment and DNA rewrapping from 1% sodium deoxycholate (aq).

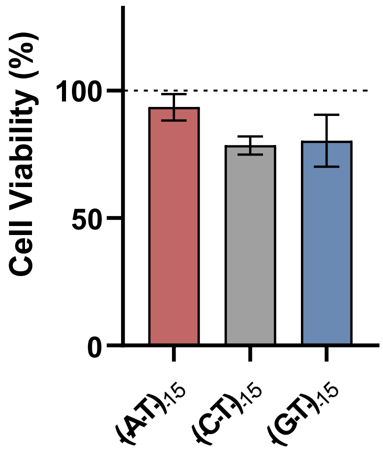

**Figure S3.** Viability of RAW 264.7 cells at DNA-SWCNT concentration of 1 mg $L^{-1}$, measured via the Tali image-based cytometry kit, after 24 hr. of incubation. N=3. Cell Viability is normalized to Control cell viability. Data are presented as mean values with error bars representing standard deviation. The experiment was performed at least three times with comparable results.

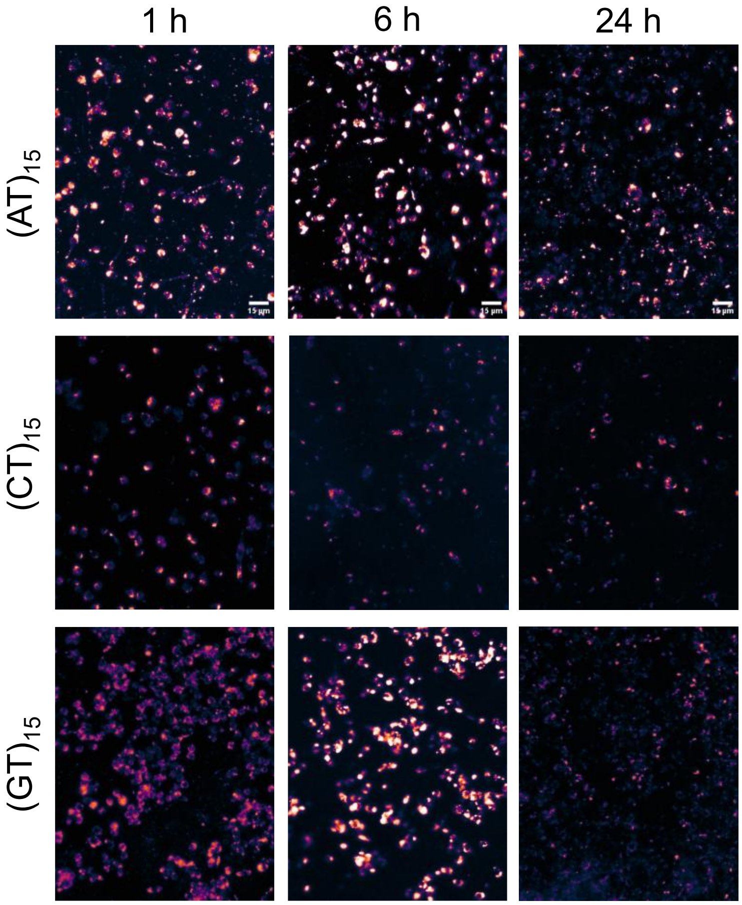

**Figure S4.** Near-infrared broadband emission of the internalized DNA-SWCNT complexes from individual puncta of the live cells at 730 nm excitation. Scale bar is 15 μm.

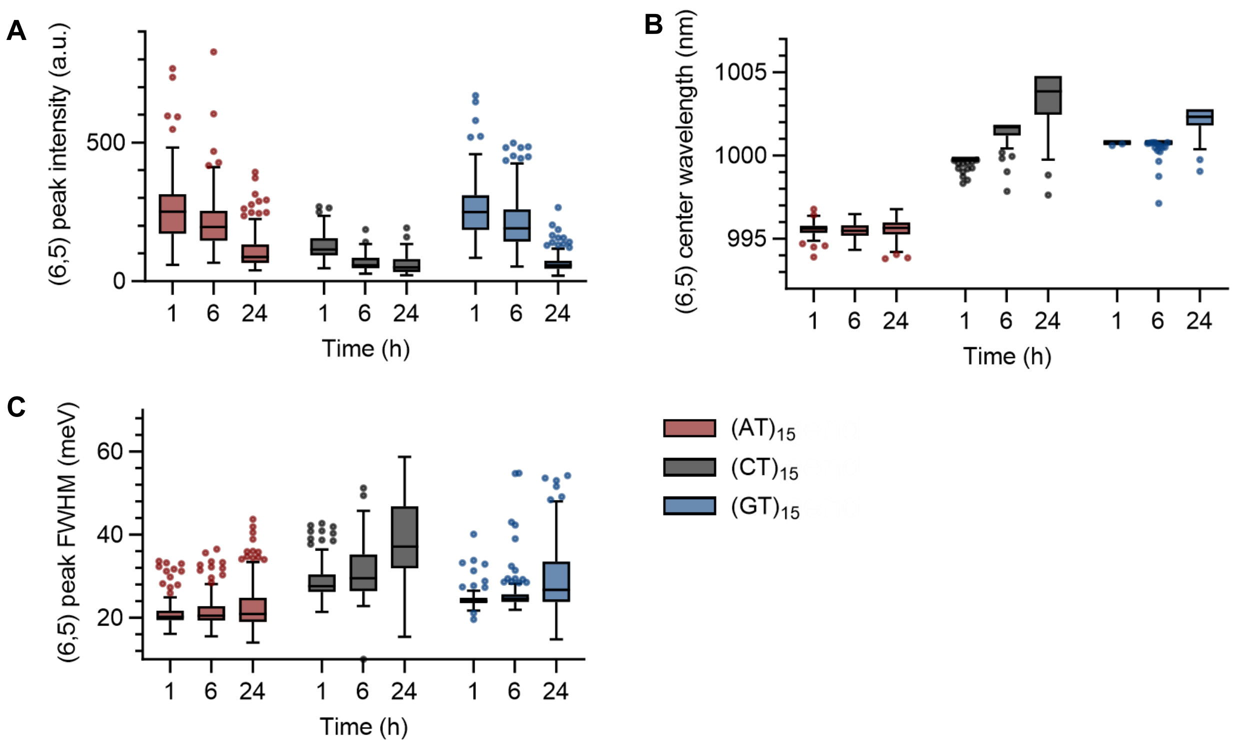

**Figure S5.** Temporal evolution of the intracellular (6,5) emission (**A**) intensity, (**B**) center wavelength, and (**C**) FWHM for each DNA-SWCNT construct over 24 hours. Boxes show the median and interquartile range (IQR), whiskers extend to 1.5x IQR (Tukey), and points beyond the whiskers are outliers. Red: $(AT)_{15}$, Black: $(CT)_{15}$, Blue: $(GT)_{15}$.

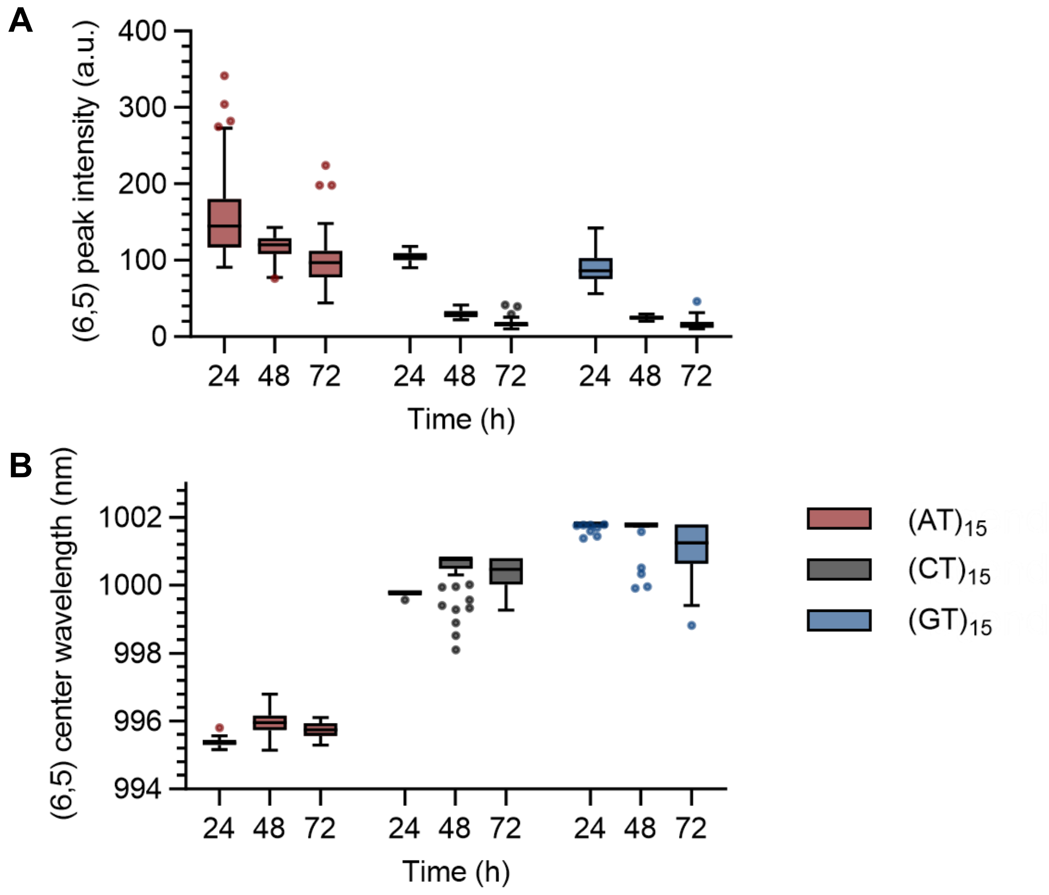

**Figure S6.** Temporal evolution of the intracellular (6,5) emission (**A**) intensity and (**B**) center wavelength for each DNA-SWCNT construct over 72 hours. Boxes show the median and interquartile range (IQR), whiskers extend to 1.5x IQR (Tukey), and points beyond the whiskers are outliers. Red: $(AT)_{15}$, Black: $(CT)_{15}$, Blue: $(GT)_{15}$.

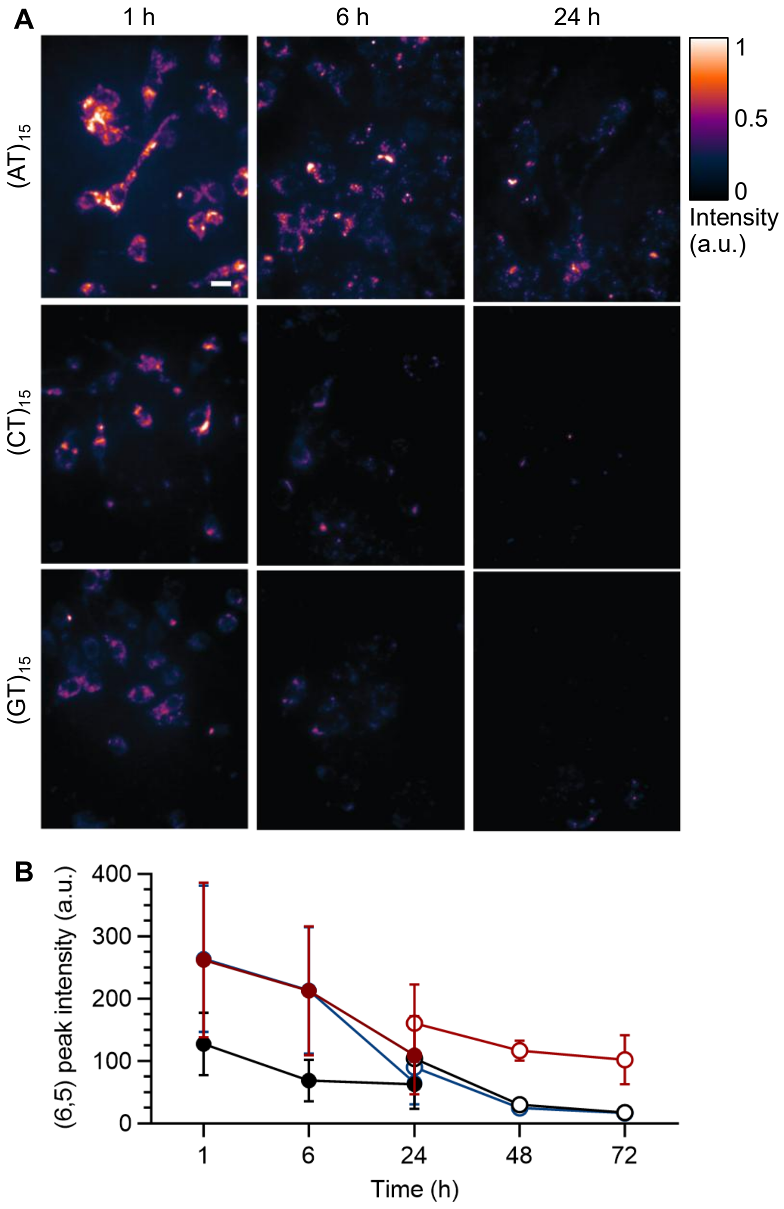

**Figure S7.** Full 72-h temporal (6,5) fluorescence intensity profiles for all three DNA-SWCNTs. (**A**) Near-infrared broadband emission of the DNA-SWCNT complexes from individual puncta of the live cells at 730 nm excitation. Scale bar: 5µm. (**B**) Temporal evolution of (6,5) intensity from 1 h to 72 h. Data represent mean ± s.d. from 3+ independent experiments. Open and filled circles denote initial DNA-SWCNT concentrations of 2 and 1 mg $L^{-1}$ for incubation, respectively. Red: $(AT)_{15}$, Black: $(CT)_{15}$, Blue: $(GT)_{15}$

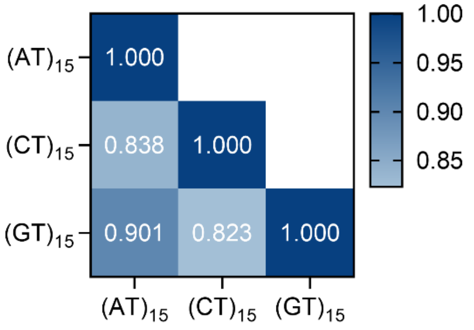

**Figure S8.** Pairwise Pearson correlation of the complete 241-protein $\log_2$ fold-change against full plasma. Color bar: correlation coefficient.

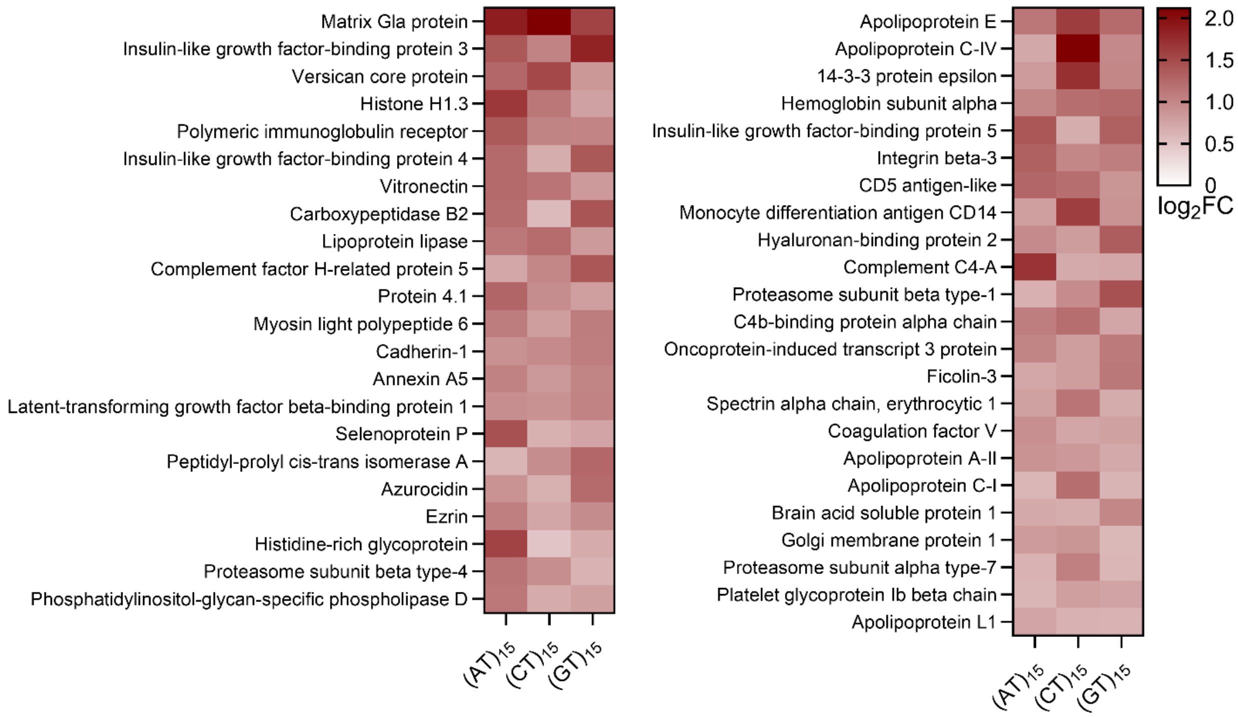

**Figure S9.** The wrapping-independent corona core. All 45 proteins enriched above 1.5-fold on all three wrappings, ordered by mean enrichment. Color encodes $\log_2$FC against full plasma. Matrix Gla protein, insulin-like growth factor binding protein 3, versican, vitronectin, and selenoprotein P are the most strongly enriched members.

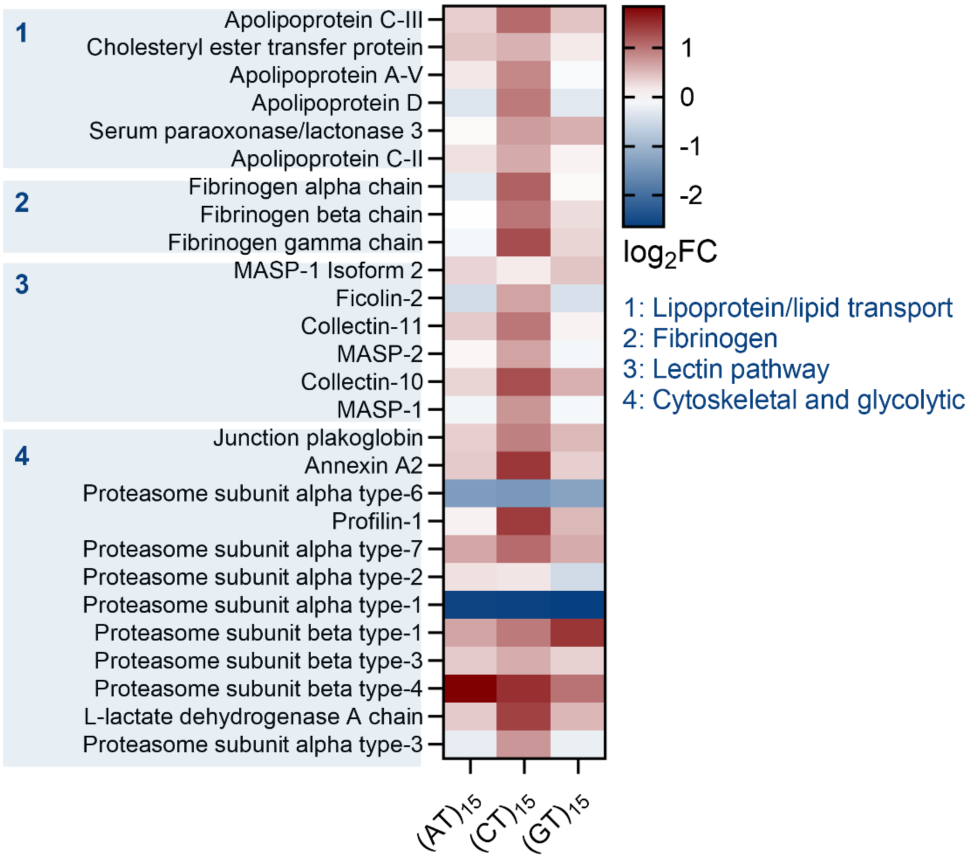

**Figure S10.** $(CT)_{15}$-SWCNT dominated corona block, grouped by functional module. 1: lipoprotein and lipid transport, 2: fibrinogen complex, 3: lectin pathway, 4: cytoskeletal and glycolytic proteins. Color encodes $log_2FC$ against full plasma.

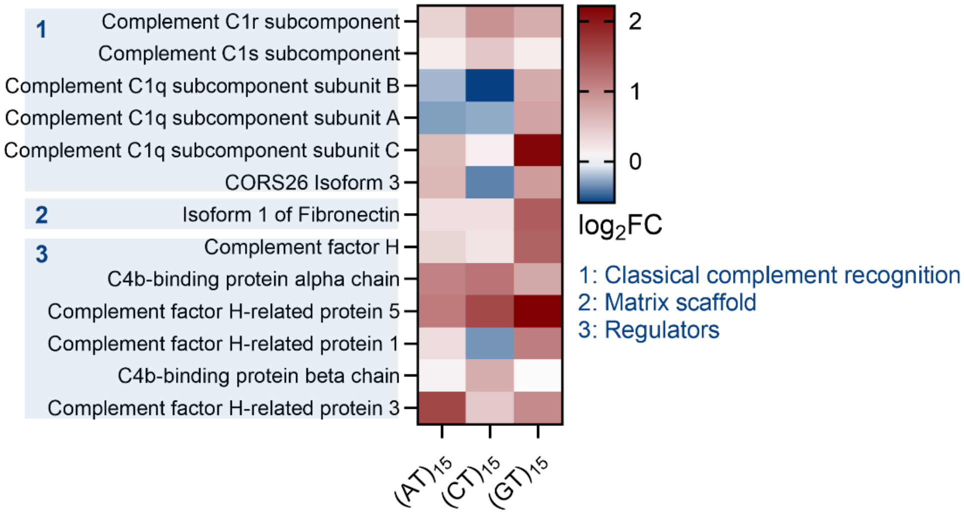

**Figure S11.** $(GT)_{15}$-SWCNT dominated corona block, grouped by functional module. 1: Classical complement recognition, 2: matrix scaffold, 3: complement regulators. Color encodes $log_2FC$ against full plasma.

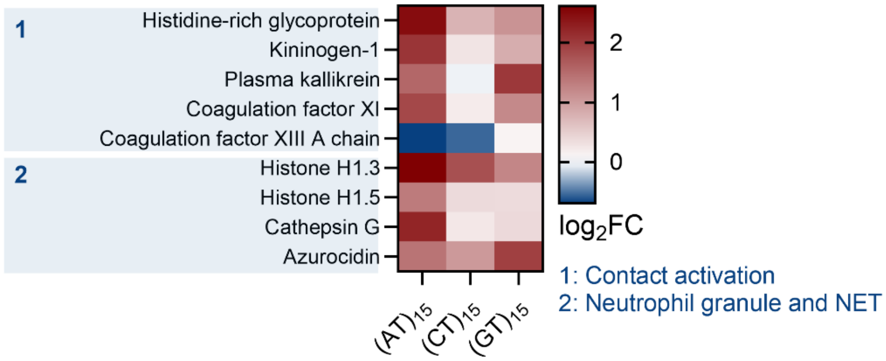

**Figure S12.** $(AT)_{15}$-SWCNT dominated corona block, grouped by functional module. 1: contact activation, 2: neutrophil granule and NET-associated proteins. Color encodes $log_2FC$ against full plasma.

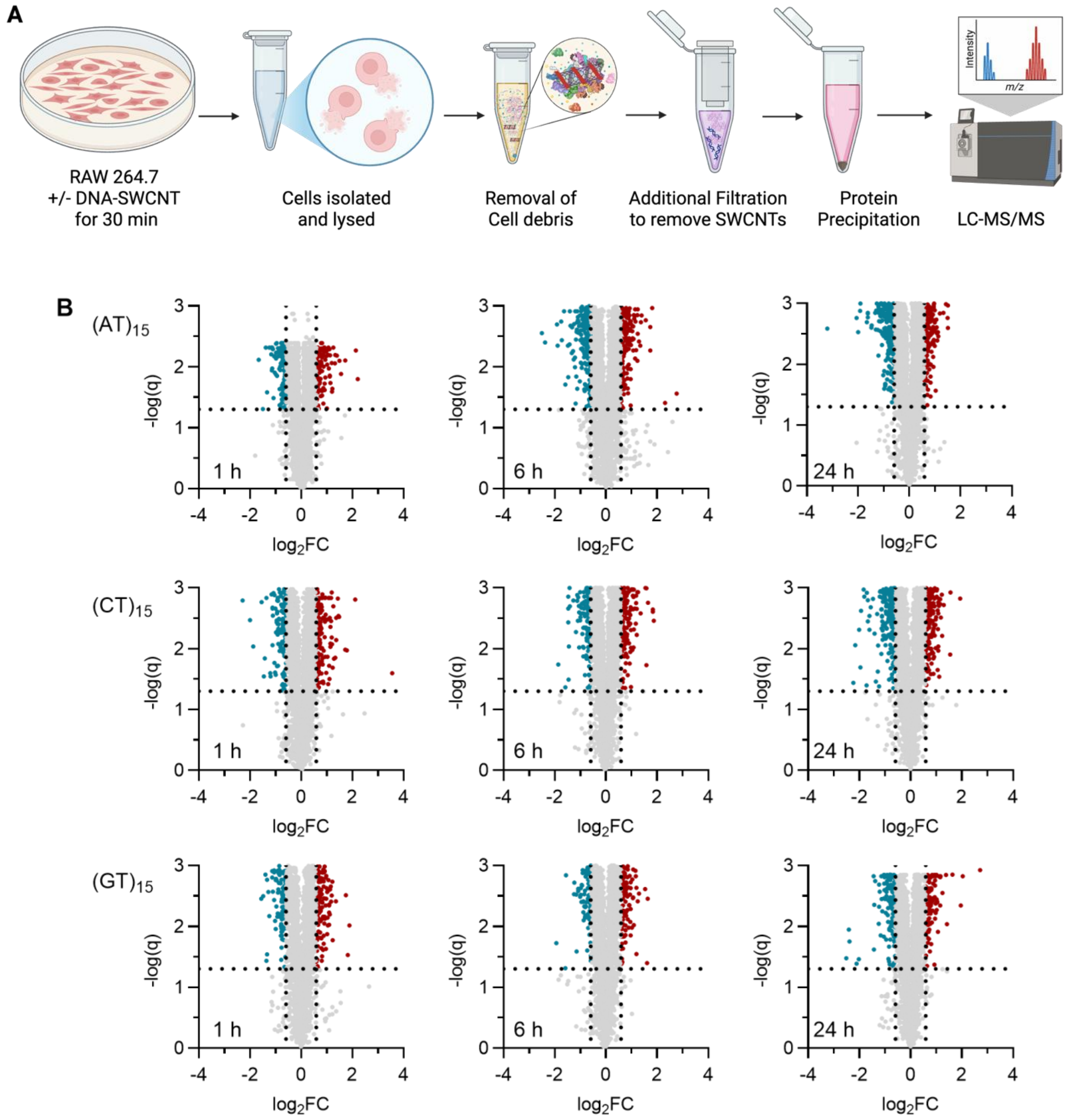

**Figure S13.** Proteomic analysis of cell lysates. (**A**) Intracellular proteomics experimental workflow. Schematic of the macrophage SWCNT exposure, lysis, TMT labeling, and LC-MS/MS pipeline used to quantify the intracellular proteome at 1, 6, and 24 h. (**B**) Volcano analysis across all conditions and time points. Volcano plots of $\log_2$ fold-change (+DNA-SWCNT vs untreated control) versus q-value (adj. p-val after BH-FDR correction) for $(AT)_{15}$, $(CT)_{15}$, and $(GT)_{15}$ at 1, 6, and 24 h. Red: Significantly enriched ($q<0.05$; $\log_2FC > 0.585$). Red: Significantly depleted ($q<0.05$; $\log_2FC < 0.585$). Gray: n.s.

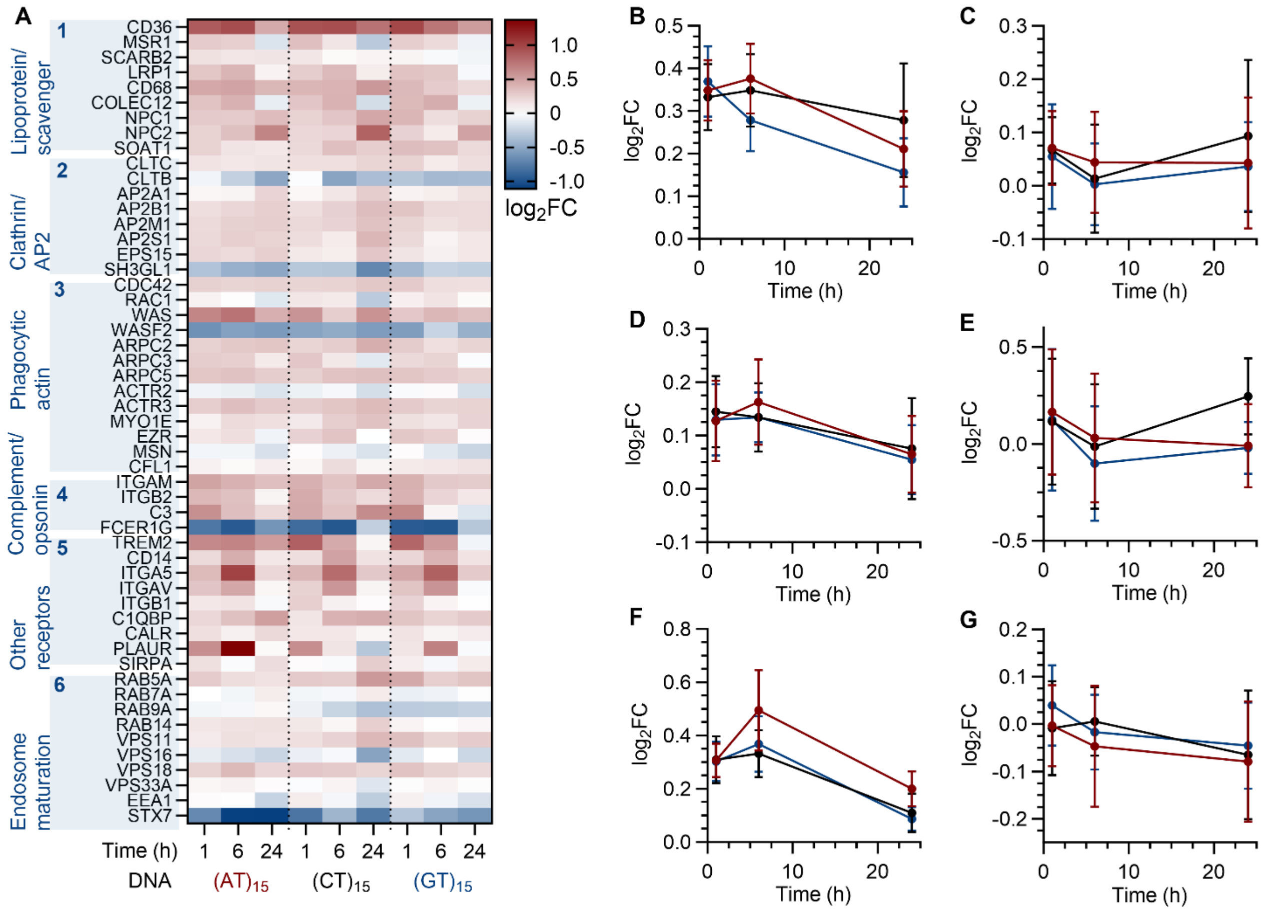

**Figure S14.** Macrophage recognition and uptake machinery is shared across the three sequences. (**A**) $Log_2FC$ against time-matched untreated control for all quantified members of six modules, 1: lipoprotein and scavenger receptors, 2: clathrin and AP2, 3: phagocytic actin, 4: complement and opsonin receptors, 5: other receptors, 6: endosome maturation. (**B**-**G**) Mean $log_2FC$ over time for each module (1-6). Red: $(AT)_{15}$, Black: $(CT)_{15}$, Blue: $(GT)_{15}$. Receptor and uptake machinery were elevated in all three constructs at comparable magnitude, with no ordering by sequence.

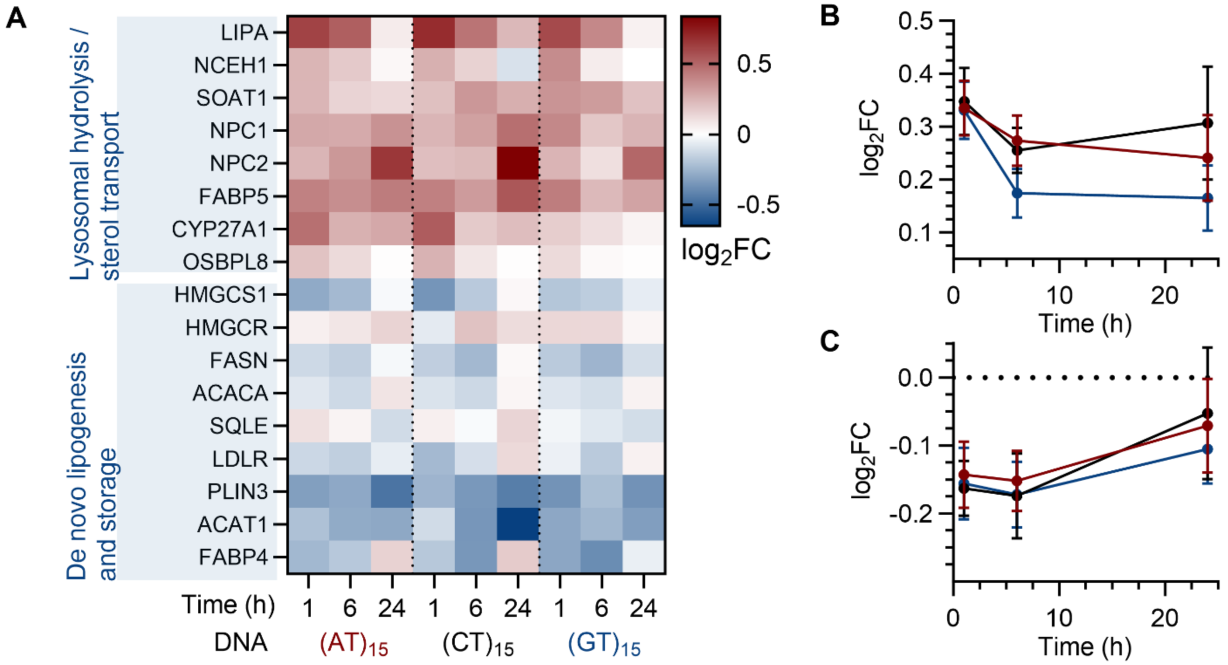

**Figure S15.** Lipid handling is engaged but not sequence specific. (**A**) $Log_2FC$ for all quantified members of two modules, 1: lysosomal hydrolysis and sterol transport, and 2: de novo lipogenesis and storage. Mean $log_2FC$ over time for (**B**) module 1 and (**C**) module 2. Red: $(AT)_{15}$, Black: $(CT)_{15}$, Blue: $(GT)_{15}$. Lysosomal lipid hydrolysis and sterol transport rose in all three constructs while lipogenesis was suppressed, and the construct with the most lipoprotein-rich corona produced no distinct response.

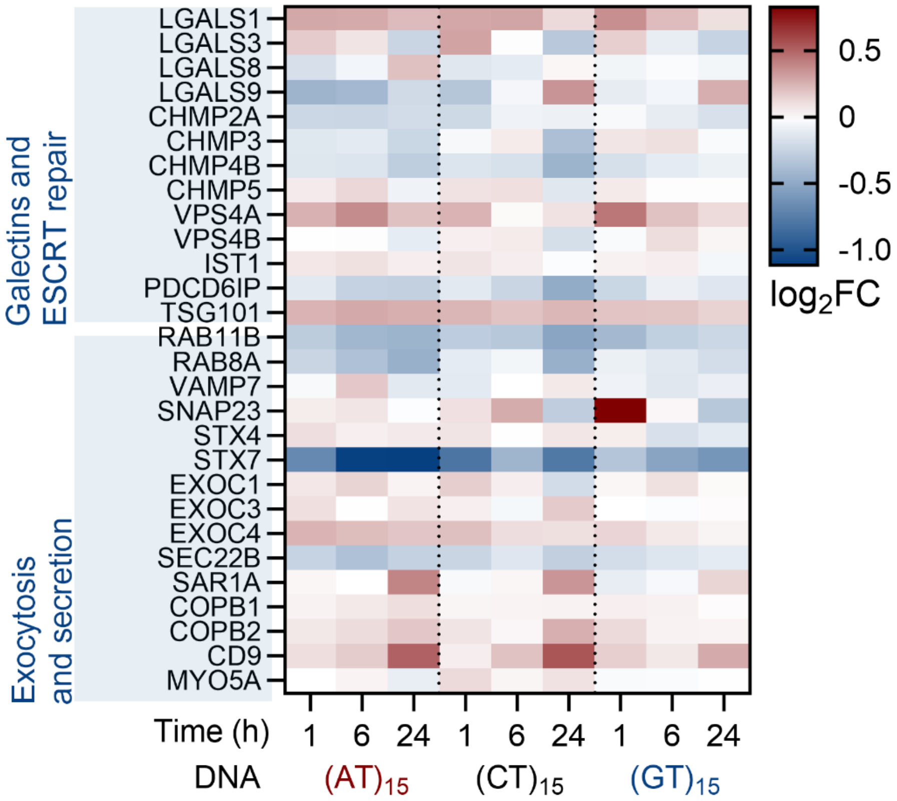

**Figure S16.** Lysosomal membrane integrity is maintained and exocytosis is not enhanced. Top block: Galectin-1, -3, -8, and -9, which accumulate on ruptured endolysosomal membranes, together with the ESCRT membrane repair proteins. Bottom block: Exocytosis and secretory machinery. No module was enriched above background in any condition.

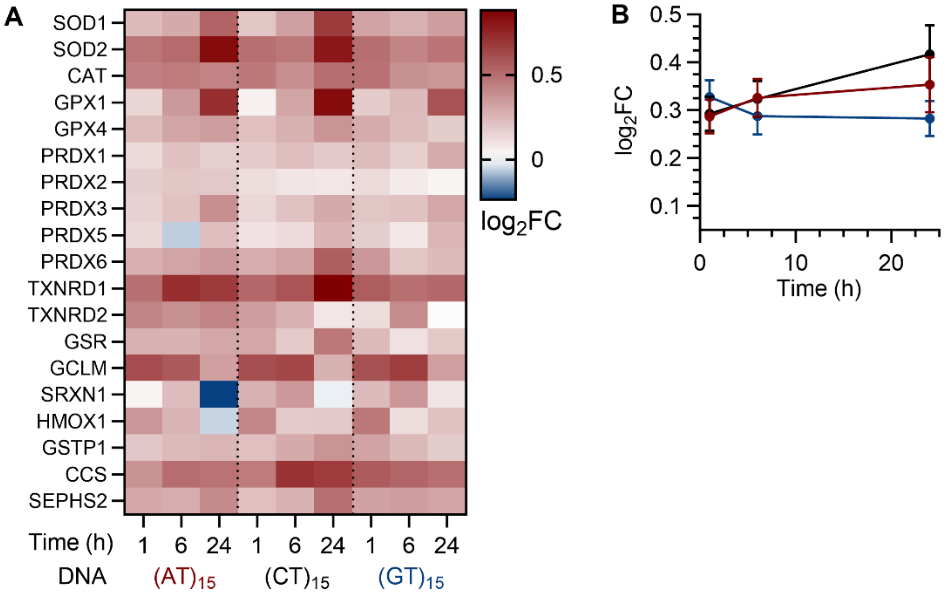

**Figure S17.** Antioxidant defenses and ROS detoxification are induced in all three sequences at every time point. (**A**) $Log_2FC$ for all quantified members. (**B**) The mean panel response over time. Red: $(AT)_{15}$, Black: $(CT)_{15}$, Blue: $(GT)_{15}$

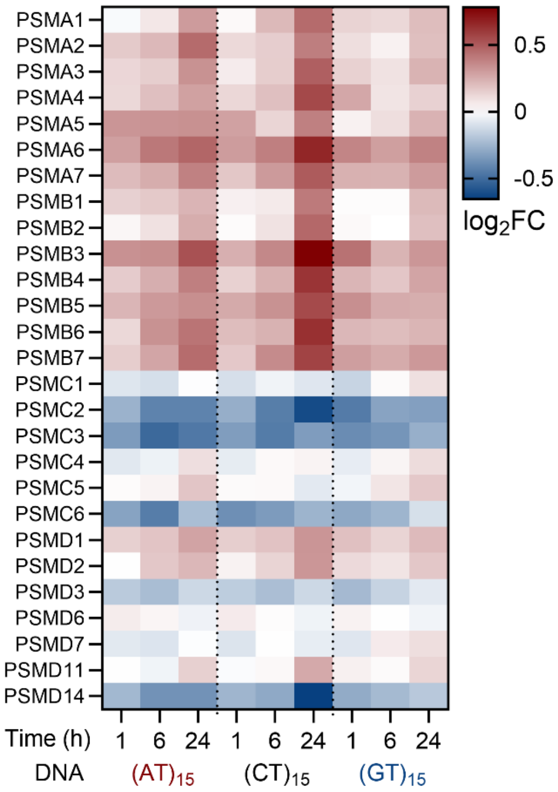

**Figure S18.** Proteasome subunits are induced at 24 h in all three sequences. $Log_2FC$ for all 27 quantified subunits.

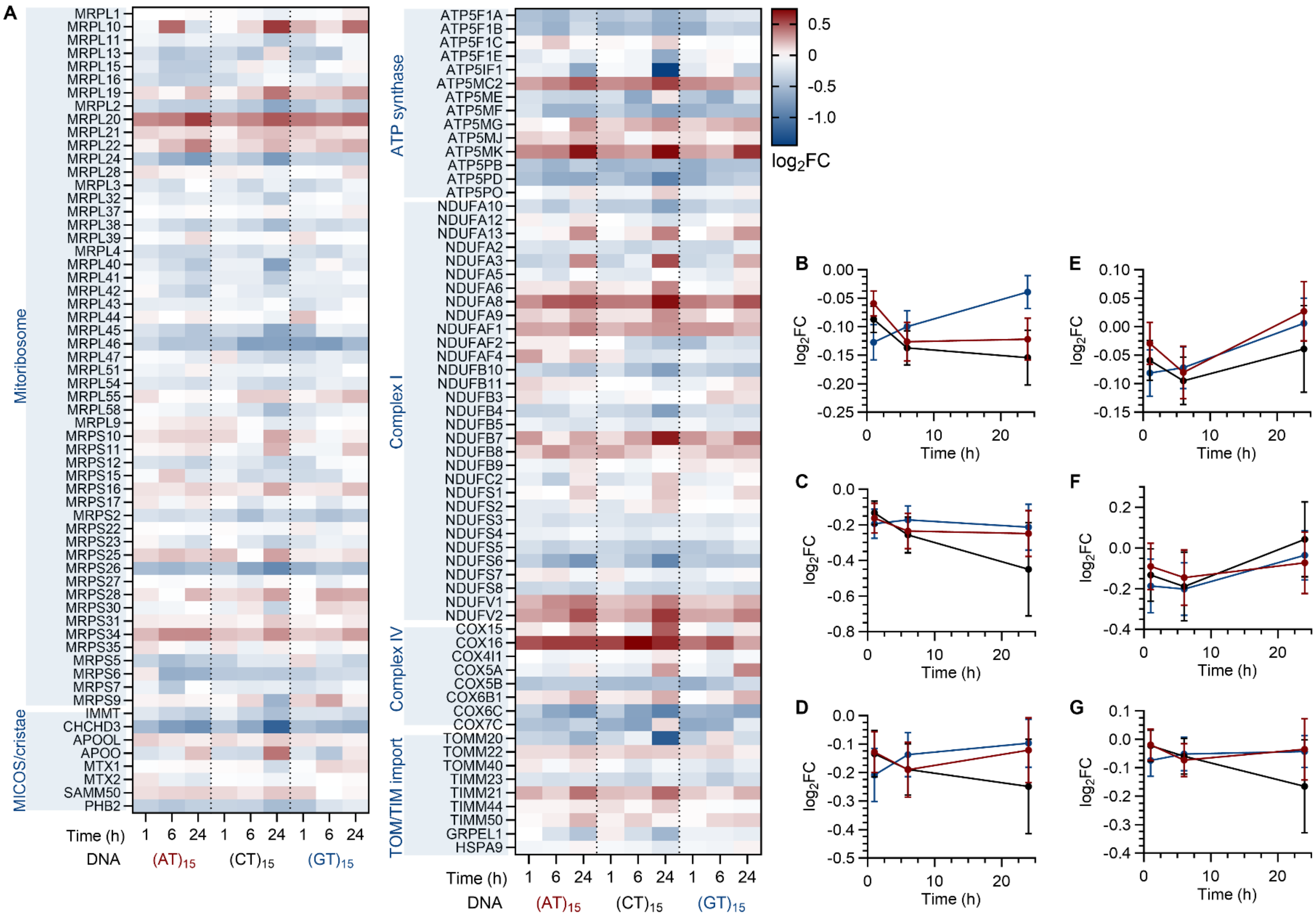

**Figure S19.** Mitochondrial proteins are depleted, and depletion is deepest in $(CT)_{15}$-SWCNT at 24 h. (**A**) $Log_2FC$ for all quantified members of the mitoribosome and the MICOS and cristae scaffold. (**B-G**) Mean $log_2FC$ over time for the mitoribosome, MICOS and cristae, ATP synthase, Complex I, Complex IV, and TOM/TIM import modules. Red: $(AT)_{15}$, Black: $(CT)_{15}$, Blue: $(GT)_{15}$.

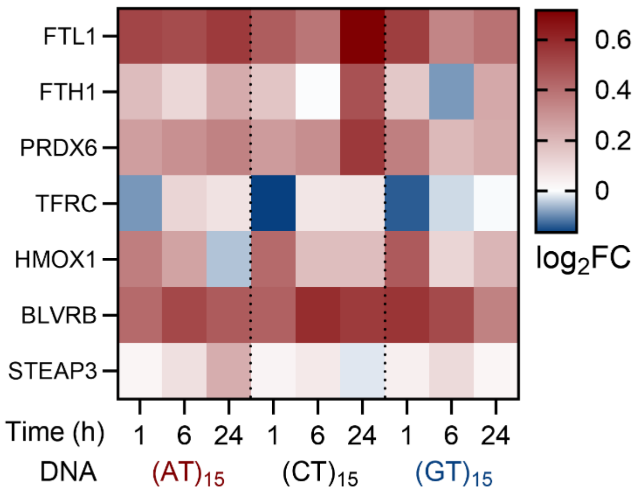

**Figure S20.** Iron and heme handling follows the same ordering as lysosomal engagement. Ferritin light chain and peroxiredoxin 6 were elevated across the time course and reached their highest values in $(CT)_{15}$-SWCNT at 24 h.
